# Elamipretide reverses female fertility decline during reproductive aging via regulating VEGF in oocytes

**DOI:** 10.64898/2026.05.27.728315

**Authors:** Hao-Lin Zhang, Yue Wang, Caizhu Wang, Xin Guo, Huanhua Chen, Yu-Xuan Hou, Xuan Wu, Zi-Jian Wu, Wen-Lin Pan, Rui-Jie Ma, Ping-Shuang Lu, Jinhui Shu, Shao-Chen Sun

## Abstract

Aging is one of the primary drivers for the decline of female fertility, and oocyte quality is the main cause for ovary aging, which is related with infertility. Although some effective anti-aging natural components have been reported, highly efficient strategies for reversing ovary aging remain lacking. In this study, we reported that peptide elamipretide reserved ovary function for female fertility during maternal aging. Our findings demonstrated that elamipretide improved aged human oocyte maturation and fertilization. Elamipretide injection increased the litter size of aged mice, and improved oocyte quality with the reverse of follicle and embryo development defect. Metabolomic and transcriptomic analyses demonstrated that multiple biological processes in oocytes were significantly reserved. Both nuclear maturation and cytoplasmic maturation of aged oocytes were improved, showing with enhancing cytoskeletal dynamics, mitochondrial metabolism and organelle rearrangement. *In vitro* supplementation during culture also restored oocyte developmental competence in both mouse and porcine oocytes. Mechanistic analysis suggested that elamipretide reversed age-related ovarian damage via synergistic activation of the Vitamin B6-VEGF axis. Therefore, our study proposed a new peptide therapy for aging-induced infertility, showing that elamipretide reverses aged oocyte quality for fertility by promoting both nuclear and cytoplasmic maturation through the coordination with VEGF signaling pathway.

## Introduction

Advanced maternal age (AMA) is associated with a variety of adverse pregnancy outcomes, including miscarriage, preterm birth, fetal growth restriction (FGR), and chromosomal abnormalities ^1–3^. Women aged 35 to 39 are more prone to infertility than those in other age groups ^4,5^. Age-related decline in oocyte quality is a key indicator of ovarian aging and an important cause of reduced fertility in elderly women. Oocyte maturation is a key process that determines the quality of oocytes, and it is complex and precisely regulated ^6^. The maturation of the oocytes includes two aspects: nuclear maturation and cytoplasmic maturation. The nuclear maturation of oocytes mainly refers to the separation of chromosomes, which reflects the ability of oocytes to resume meiosis; and the cytoplasmic maturation of the oocytes involves the accumulation of mRNAs, proteins, and substrates required for subsequent fertilization/development^7,8^. The cytoplasmic maturation failure induced by aging is associated with mitochondrial dysfunction, alterations in mitochondrial biogenesis, changes in mitochondrial morphology, distribution, activity and dynamics, smooth endoplasmic reticulum malformation, calcium disturbance, and cytoskeletal changes ^9^. Previous studies have found oxidative damage in oocytes of AMA and confirmed that these oocytes have mitochondrial dysfunction ^10^. And mitochondria dysfunction is an important reason led to decreased oocyte quality in aged mice or females ^11^. Besides, age-related changes in polar metabolites suggested a decrease in mitochondrial function, as demonstrated by NAD+, purine, and pyrimidine depletion, while glycolysis substrates and glutamine accumulated, with age ^12^. Mitochondria are the multifunctional powerhouse of cells, including human oocytes, by producing oxidative phosphorylated ATP (OXPHOS) through five complexes (Complex I-V) in the electron transport chaint^13^. Thus, mitochondria are key organelles that provide energy for oocyte maturation, fertilization, and embryonic development. In aging ovaries or oocytes, the main feature of mitochondrial dysfunction is the decline in the respiratory capacity of each mitochondrial, while the mitochondrial membrane potential (MMP) drops in a stable state ^14^. In aging cells, relatively low MMP is typically associated with increased production of reactive oxygen species (ROS) which triggers oxidative stress, in turn leads to nuclear DNA and mitochondrial damage, insufficient intracellular energy supply, calcium homeostasis imbalance, and meiotic spindle abnormalities, ultimately potentially resulting in oocyte aneuploidy^15–17^.

Multiple studies focused on investigating targeted rescue strategies, including regulating endogenous metabolism, supplementing natural extracts, and applying small-molecule drug inhibitors, for improving the quality of aged oocytes. The small molecule 78c, as an inhibitor of CD38 (ADP-ribosyl cyclase/hydrolase), was demonstrated to elevate NAD^+^ levels and reduce inflammation in aging ovaries, thereby improving oocyte quality ^18^. While the generation of young follicle-aged oocyte chimeras improved the developmental quality of embryos following *in vitro* fertilization and embryo implantation ^19^. Besides, regulation of endogenous metabolism such as mevalonate (MVA) and supplementation of natural extract like 8-isopentenyl flavone provides (8-IPF) could promote the synthesis of FPP in adjacent granulosa cells, thereby restoring the assembly of cortical actin in aged oocytes ^20^. In addition, there are relevant studies on using extracts to rescue the quality of aged oocytes. Rapamycin was proven to ameliorate age-related infertility caused by ribosomal dysfunction via restoring protein homeostasis ^21^. It was found that melatonin eliminated oxidative stress-induced meiotic defects and aneuploidy in aged mouse oocytes ^22^. Elevating spermidine levels could improve follicular development, oocyte maturation, early embryonic development, as well as female fertility in aged mice ^23^. Additionally, antioxidant interventions specifically directed towards mitochondria represent a key approach in this field. MitoQ and SkQ1, which could effectively reduce ROS generation and preserves mitochondrial structure and function through mitochondria-targeted antioxidant properties ^24,25^. Nicotinamide mononucleotide (NMN) and NAD+ enhancers, which could promote cellular metabolism, energy production, and mitochondrial function by increasing the intracellular NAD+ levels ^26^. CoQ10 mainly functions in the mitochondrial electron transport chain, serving as an electron carrier to transfer electrons from complexes I and II to complex III, driving the proton pump, helping to establish a membrane electrochemical gradient, and promoting ATP synthesis ^27^. However, these therapeutic strategies still suffer from drawbacks such as low targeting efficiency and poor bioavailability. Therefore, it is imperative to explore safer and more readily absorbable small-molecule agents for clinical translation.

Arg-Dmt-Lys-Phe-NH2 (also known as MTP-131, elamipretide, and Bendavia, collectively referred to as elamipretide) is a cell membrane-penetrating aromatic cationic tetrapeptide with a molecular weight of 639.8 g/mol, composed of four amino acids with both hydrophilic and hydrophobic properties^28^. It is reported that elamipretide can improve mitochondrial function and reduce oxidative stress damage ^29^. Combining the advantages of elamipretide such as small molecular size, easy synthesis, good water solubility, not easily degraded by peptidase, and high stability in solution, elamipretide is considered to be a potential mitochondrial-targeted drug that can be applied in clinical practice. Under physiological pH conditions, elamipretide carries three positive charges and selectively targets and accumulates in the inner mitochondrial membrane through electrostatic and hydrophobic interactions ^30^. It has been demonstrated to bind cardiolipin with high affinity. In ischemia reperfusion injury, it formed a complex with cardiolipin on the inner mitochondrial membrane, thereby inhibiting cytochrome *c* peroxidase activity and protecting mitochondrial cristae ^31^. Cardiolipins in mitochondria were prone to oxidation under oxidative stress, resulting in membrane disruption, reduced electron transport chain (ETC) efficiency, and diminished ATP production ^32^. The specific binding of elamipretide to cardiolipin effectively prevented cardiolipin oxidation ^33^. elamipretide thus stabilized the integrity of mitochondrial membranes and enhanced the efficiency of the electron transport chain (ETC), thereby optimizing the overall bioenergetic function of mitochondria ^34^. elamipretide could improve mitochondrial structure and function, thereby further reducing the production of reactive oxygen species (ROS) in mitochondria and minimizing the damage of oxidative stress to mitochondria ^35^. Additionally, in human retinal endothelial cells, elamipretide was found that it could also lower abnormal mitochondrial membrane potential, prevented excessive calcium ions from entering mitochondria, and inhibited cell apoptosis ^36^. These effects of elamipretide collectively enhance mitochondrial energy metabolism efficiency, stabilize ATP production, and reduce oxidative stress damage ^37–39^. Report indicated that elamipretide treatment is effective at mitigating signs of sarcopenia and cardiac dysfunction in an aging mouse model ^40^. Another study showed long treatment with elamipretide could improve the physical performance of aged males ^41^.

In this study, we tried to explore whether the treatment of elamipretide could reverse the fertility decline of aging females and the potential mechanism. We found that elamipretide synchronously promotes oocyte nuclear-cytoplasmic maturation by activating the VB6-VEGF axis, and enhances embryo quality to reverse age-related fertility decline in mice.

## Materials and Method

### Patients

This study was approved by the Ethics Committee of Guangxi Maternal and Child Health Hospital and conducted in accordance with the Declaration of Helsinki. Written informed consent was obtained from each participant. All resulting oocytes were used solely for research purposes to assess oocyte developmental competence and were not transferred. Patients undergoing intracytoplasmic sperm injection (ICSI) cycles were enrolled. The inclusion criteria were: 1) meeting the clinic’s indications for ICSI, and 2) at least two germinal vesicle (GV) oocytes retrieved. The exclusion criterion was a maturation rate below 60%.

### Randomization and Procedures

Ovarian stimulation was performed using the antagonist protocol. In brief, recombinant FSH was initiated on menstrual cycle days 2–3. A GnRH antagonist (Cetrotide, MSD China) was introduced when follicles reached 11–13 mm and luteinizing hormone (LH) ≥10 IU/L or estradiol (E2) ≥1570 pmol/L. Human chorionic gonadotropin (hCG, 6000–8000 IU) was administered when at least three follicles reached ≥17 mm or at least two follicles reached ≥18 mm. Oocytes were collected by transvaginal ultrasound□guided follicular aspiration 34–36 hours after hCG administration. Cumulus□oocyte complexes (COCs) were promptly transferred to the incubator in commercial cleavage culture medium (Vita, Shenzhen, China). After 3 hours of incubation, COCs were denuded using hyaluronidase to remove cumulus cells, and metaphase II (MII) oocytes were selected for ICSI. GV oocytes were collected for in vitro maturation (IVM).

All cultures were performed in a humidified tri□gas incubator at 37°C with 89% N□, 6% CO□, and 5% O□. GV oocytes were divided into two groups based on patient age: an advanced-age group (≥35 years) and a younger group (<35 years). Within each age group, GV oocytes were randomly allocated to either the experimental group or the control group. Experimental group: commercial IVM medium supplemented with 500 μM SS□31 peptide; Control group: commercial IVM medium alone. The maturation media were prepared 4–6 hours in advance and equilibrated at 37°C in 89% N□, 5% O□. Both groups were cultured in the same incubator under 37°C in 89% N□, 5% O□ for observation. Oocytes that extruded the first polar body were defined as mature MII oocytes. Mature oocytes (MII) obtained at 24 hours were subjected to ICSI using surplus sperm sample donated for research purposes, and this day was designated as Day 0.

Fertilization was assessed 16–18 hours after ICSI by the presence of two pronuclei (2PN). Normally fertilized zygotes were cultured individually in sequential medium (Vita, Shenzhen, China). Embryos were morphologically evaluated on day 3 (67–69 hours post□insemination) and then cultured to the blastocyst stage in blastocyst medium (Vita, Shenzhen, China). Day 3 embryos with a grading score ≥6C□II and fragmentation≤20% were classified as good quality embryos.

### Outcome Measures

The following outcomes were assessed: maturation rate at 24 hours, maturation rate at 48 hours, fertilization rate, cleavage rate, good quality embryo rate, and blastocyst formation rate.

### Antibodies and chemicals

Mouse monoclonal anti-α-tubulin-FITC antibody was from Sigma-Aldrich Corp (St. Louis, MO, USA, Cat# F-2168-2ML, 1:400). TRITC-Phallodin was purchased from SAITONG (Beijing Pusitang Biotechnology Co., LTD, Cat# T10446-300 T, 1:200). Rabbit anti-gamma tubulin antibody, and rabbit anti-Rab10 antibody were from abcam (Cambridge, UK, Cat# ab179503, 1:200). Mito-Tracker Red CMRos (1:500, Cat# M7521, Invitrogen, Eugene, OR, USA) was used to detect the distribution of mitochondria with an ultimate density of 2μmol/L. TMRE (Cat# C2001S, 1:200) was used to detect the mitochondrial membrane potential. ROS (Cat# S0033S, 1:800), Annexin-V (Cat# C1062S, 1:10), Golgi-tracker Green (Cat# C1045S-1, 1:100), ER-tracker Green (Cat# C1042M-1, 1:100), and lysosome-tracker Red (1:12000, Cat# C1046) were purchased from Beyotime Biotechnology (Nantong, China). Fluo-4 AM was purchased from Beyotime Biotechnology (Nantong, China). Rabbit anti-LC3B antibody was purchased from Cell Signal Technology (Cat #2775, 1:100). PARK2/Parkin Polyclonal antibody (Cat# 14060-1-AP, 1:200) and GRP78/BIP (11587-1-AP) was from Proteintech Group, Inc (Rosemont, IL, USA). Lens Culinaris Agglutinin (LCA)-Fluorescein (FITC) was from Invitrogen Corporation (Carlsbad, CA, USA, Cat# L32475). Rabbit monoclonal anti-VEGF, anti-PI3K and anti-AKT2 was purchased from Cell Signal Technology. Beta-actin and α-tubulin were purchased from Cell Signal Technology. Goat anti-rabbit IgG/Alexa Fluor 488 (Cat# ZF-0511, 1:200) and TRITC-conjugated goat anti-rabbit IgG (Cat# ZF-0316, 1:200) were from Zhongshan Golden Bridge Biotechnology, Co., Ltd. (Beijing, China). Horseradish peroxidase-conjugated goat anti-rabbit/mouse IgG (H + L) antibodies (Cat# CW0102S, 1:2000) were obtained from CWBIO (Beijing, China). Emvododstat (PTC299) (E1078), LY294002(S1105), and Wortmannin (S2758) was purchased from Selleckchem.com. All other chemicals were purchased from Sigma (St. Louis, MO, USA), unless otherwise stated.

### Mouse oocyte collection and culture

All mouse experiments were meticulously adhered to the protocols established by the Animal Research Institution of Nanjing Agricultural University in China, and the experimental procedures were formally approved by the Experimental Animal Research Committee. In the in vitro experiments, germinal vesicle (GV)-stage oocytes were meticulously harvested from the ovaries of both 10-month-old female mice and 6- to 8-week-old ICR female mice. Following a thorough cleansing procedure, these oocytes were subsequently transferred into M2 culture medium, overlaid with a layer of paraffin oil to maintain a stable microenvironment. The oocytes were then cultured under precisely controlled conditions of 37□ and 5% CO□ for durations of 8 and 12 hours, respectively. This protocol enabled the successful acquisition of oocytes at the metaphase I (MI) and metaphase II (MII) stages, which were subsequently utilized for further experimental procedures.

### Porcine oocyte collection and IVM (*in vitro* maturation)

All procedures followed the Animal Research Protocol approved by Nanjing Agricultural University. Ovaries from prepubertal gilts were recovered at a local abattoir, held in 0.9 % saline at 38 °C, and delivered to the laboratory within 2 h. Following two DPBS washes, 3–6 mm antral follicles were punctured with 20-gauge needles; only COCs showing intact, compact cumulus investment and homogeneous ooplasm were kept. Basal maturation medium was TCM-199 fortified with 75 µg mL□¹ penicillin, 50 µg mL□¹ streptomycin, 0.5 µg mL□¹ FSH, 0.5 µg mL□¹ LH, 10 ng mL□¹ EGF and 0.57 mM cysteine. Groups of 25–30 COCs were matured in 500 µL droplets under mineral oil at 38.5 °C, 5 % CO□, for 27 or 44 h. Cumulus cells were then removed by a 5-min exposure to 0.1 % hyaluronidase at 38.5 °C and gentle pipetting; denuded oocytes were washed 3–4 times and examined under an inverted microscope (×200) for polar-body extrusion.

### *In vitro* fertilization and embryo culture

The animal experiments in this study were approved by the Animal Ethics Committee of Nanjing Agricultural University and were conducted in strict compliance with the guidelines of the Animal Research Committee. ICR female mice (purchased from Nanjing Medical University) at 10 months and 6–8 weeks of age were induced to superovulate through injection of 10 IU of PMSG, followed by 10 IU of hCG 46-48 hours later. ICR male mice (9-10 weeks old, purchased from Qinglongshan Animal Farm) were sacrificed 13 hours after hCG injection. Their epididymides were harvested and capacitated in prewarmed HIF medium for 1 hour at 37°C under 5% CO□. Approximately 14 hours after hCG injection, cumulus-oocyte complexes were retrieved from the ampullae of the female mice’s oviducts and placed into HTF fertilization drops. Subsequently, capacitated sperm were introduced into the fertilization drops and co-incubated with oocytes for 4-6 hours at 37°C under 5% CO□ for fertilization. Finally, the fertilized eggs were collected and cultured in KSOM medium (37°C, 5% CO□) until they developed into early embryos, such as the 2-cell and 4-cell stages.

### Elamipretide treatment

In the in vitro experiments, elamipretide powder was initially dissolved in normal saline to create a 100 mM concentrated stock solution. This stock solution was then serially diluted with M16 culture medium to yield working solutions of 100 μM, 200 μM, and 400 μM. Subsequently, GV-stage oocytes derived from both young and aged mice were transferred into the elamipretide-containing culture medium. These oocytes were cultured for 12 hours under standardized conditions of 37□ and 5% CO□ to assess oocyte maturation rates. Ultimately, a concentration of 200 μM elamipretide was determined to be the most effective for rescue purposes.

In the *in vivo* experimental design, elamipretide powder was dissolved in normal saline to formulate a 100 mM solution, which was then aliquoted and stored at -20°C. Prior to use, the solution was thawed at room temperature and administered to aged mice via intraperitoneal injection at three different dosages: 3 mg/kg/day, 5 mg/kg/day, and 10 mg/kg/day. The efficacy of these dosages was evaluated, and a dose of 5 mg/kg/day was identified as the optimal therapeutic dose. To further optimize the treatment protocol, three different treatment durations—3 days, 7 days, and 10 days—were tested. Ultimately, a 7-day treatment duration was determined to be the most effective therapeutic regimen.

### Immunofluorescent staining and confocal microscopy

At least 30 oocytes were initially fixed in a 4% paraformaldehyde solution in PBS (pH 7.4) for 30 minutes. Following fixation, the oocytes were permeabilized using 0.5% Triton-X-100 for 20 minutes at room temperature. After permeabilization, the oocytes were blocked with 1% bovine serum albumin (BSA) and subsequently incubated with primary antibodies, including γ-tubulin (1:200), acetylated tubulin (1:400), Rab10 (1:500), GRP78(1:200), LC3(1:100), and anti-α-tubulin-FITC (1:100), overnight at 4°C. After the incubation period, the oocytes were transferred to a washing solution composed of 0.1% Tween 20 and 0.01% Triton X-100 in PBS and subjected to three washing cycles. Subsequently, the oocytes were labeled with Alexa Fluor 488 or Alexa Fluor 594-conjugated secondary antibodies at a dilution of 1:100. To visualize F-actin via phalloidin staining, the oocytes were subjected to incubation with phalloidin-TRITC at room temperature for a duration of 1 hour. Subsequent to this step, the chromosomes within the oocytes were stained using Hoechst 33342 for a period of 10 minutes. The oocytes were subsequently affixed to glass slides and analyzed using a confocal laser scanning microscope fitted with a 40 × water immersion objective lens (Zeiss LSM 700 META, Germany). Moreover, live cell fluorescence staining was employed to evaluate the distribution of mitochondria with Mito Tracker Red CMXRos (Invitrogen, USA), mitochondrial membrane potential (Mito Probe JC-1 Assay kit, Invitrogen, USA), and calcium level (Fluor 4.AM, Beyotime Biotechnology, Nantong, China), Golgi (Beyotime Biotechnology, Nantong, China), endoplasmic reticulum (Beyotime Biotechnology, Nantong, China) and lysosome (Invitrogen, Eugene, OR, USA). Prior to transfer into M2 medium supplemented with fluorescent probes, oocytes were cultured to reach the desired meiotic stage. Following a 30-minute incubation at 37°C under 5% CO□ conditions, the oocytes were immediately subjected to confocal microscopy imaging.

### MtDNA copy-number assessment

Fifty oocytes per group were pooled and processed with the Bioengineering mitochondrial DNA isolation kit (B518749-0050). Relative mtDNA abundance was determined by qPCR, normalizing the mitochondrial amplicon to the nuclear β-actin sequence. Primer pairs (synthesized by Genewiz) were: β-actin: 5′-TGT GAC GTT GAC ATC CGT AA-3′ and 5′-GCT AGG AGC CAG AGC AGT AA-3′; mitochondrial target: 5′-CCA ATA CGC CCT TTA ACA AC-3′ and 5′-GCT AGT GTG AGT GAT AGG GTA G-3′.

### ATP quantification

Thirty oocytes per group were lysed and processed with the Sigma bioluminescent ATP kit (Cat# 1003493051). After brief permeabilization with the supplied, pre-diluted ATP-release reagent, samples were combined with luciferase substrate and immediately read on a Tecan Spark multimode reader; emitted light was converted to relative ATP concentrations.

### Western blotting

Around 50 mouse oocytes were solubilised in NuPAGE LDS sample buffer, heated (10 min, 100 °C), and kept at −20 °C until use. Equal volumes were loaded on 10 % SDS-polyacrylamide gels and resolved at 160 V for 70 min. Proteins were electro-transferred to PVDF membranes (Millipore, Billerica, MA) at 20 V for 90 min. After blocking with fast blocking buffer (0.5 h, RT), membranes were probed overnight at 4 °C with primary antibodies (rabbit anti-DRP1, rabbit anti-α-tubulin, mouse anti-β-actin, anti-GM130, anti-GRP78 and anti-acetyl-tubulin, anti-LC3, anti-ROCK1, anti-Park2/PRKN, all 1:1000). Following five TBST washes (10 min each), HRP-conjugated goat anti-rabbit or anti-mouse IgG (1:2000) was applied for 1 h at RT. Signals were developed with YEASEN high-sig/super-sig ECL (1:10) and quantified by ImageJ.

### Real-time quantitative PCR analysis

Real-time quantitative PCR (qPCR) was employed to assess the mRNA expression level of genes. Total RNA was extracted from a pool of 30 oocytes utilizing the Dynabeads mRNA DIRECT kit (Invitrogen Dynal AS, Norway). Following RNA isolation, first-strand cDNA synthesis was conducted using the PrimeScript RT Master Mix (Takara, Japan) with the thermal cycling conditions of 37°C for 15 minutes, 85°C for 5 seconds, and a final hold at 4°C. The primer sequences used for amplifying the cDNA fragments of the genes are listed in the table in the attachment. The PCR protocol was carried out under the following conditions: an initial denaturation at 95°C for 30 seconds, succeeded by 40 cycles of quantitative PCR (qPCR) amplification (95°C for 5 seconds and 60°C for 30 seconds), and concluded with a melt-curve analysis comprising 95°C for 5 seconds, 60°C for 60 seconds, and a final step at 95°C for 1 second, with the reaction maintained at 4°C. The primers are listed in Table S1.

### Statistical analysis

For each experimental analysis, a minimum of three biological replicates was performed to ensure reliability. The mean values and standard error of the mean (SEM) were calculated. Statistical significance between the three groups was evaluated using ordinary One-Way ANOVA. The statistical analyses were carried out within GraphPad Prism 5 software. Results were deemed statistically significant when the P-value was less than 0.05 (indicated by *) and highly significant when the P-value was less than 0.01 (indicated by **).

## Results

### Elamipretide treatment improves human oocyte maturation, fertilization and cleavage

A total of 14 patients were enrolled. In the advanced-age group (≥35 years), three patients (mean age 36.33 ± 0.58 years) contributed 27 GV oocytes, which were randomly assigned to the experimental (n = 15) or control (n = 12) group. In the younger group (<35 years), four patients (mean age 29.50 ± 2.38 years) contributed 60 GV oocytes, assigned to the experimental (n = 31) or control (n = 29) group. We first tested the effects of elamipretide on human oocytes of both aged women (>36 years) and young women. As shown in Table 1, *in vitro* culture of human aged oocytes with elamipretide improved the maturation with both 24 hours culture and 48 hours culture, while there is a significant increase for the fertilization rate and cleavage rate after ICIS with MII oocytes. Similarly, elamipretide supplement on the oocytes of young women also showed higher maturation rate and cleavage rate. These data suggested that elamipretide treatment could improve oocyte maturation and developmental competence in human.

**Table 1.**
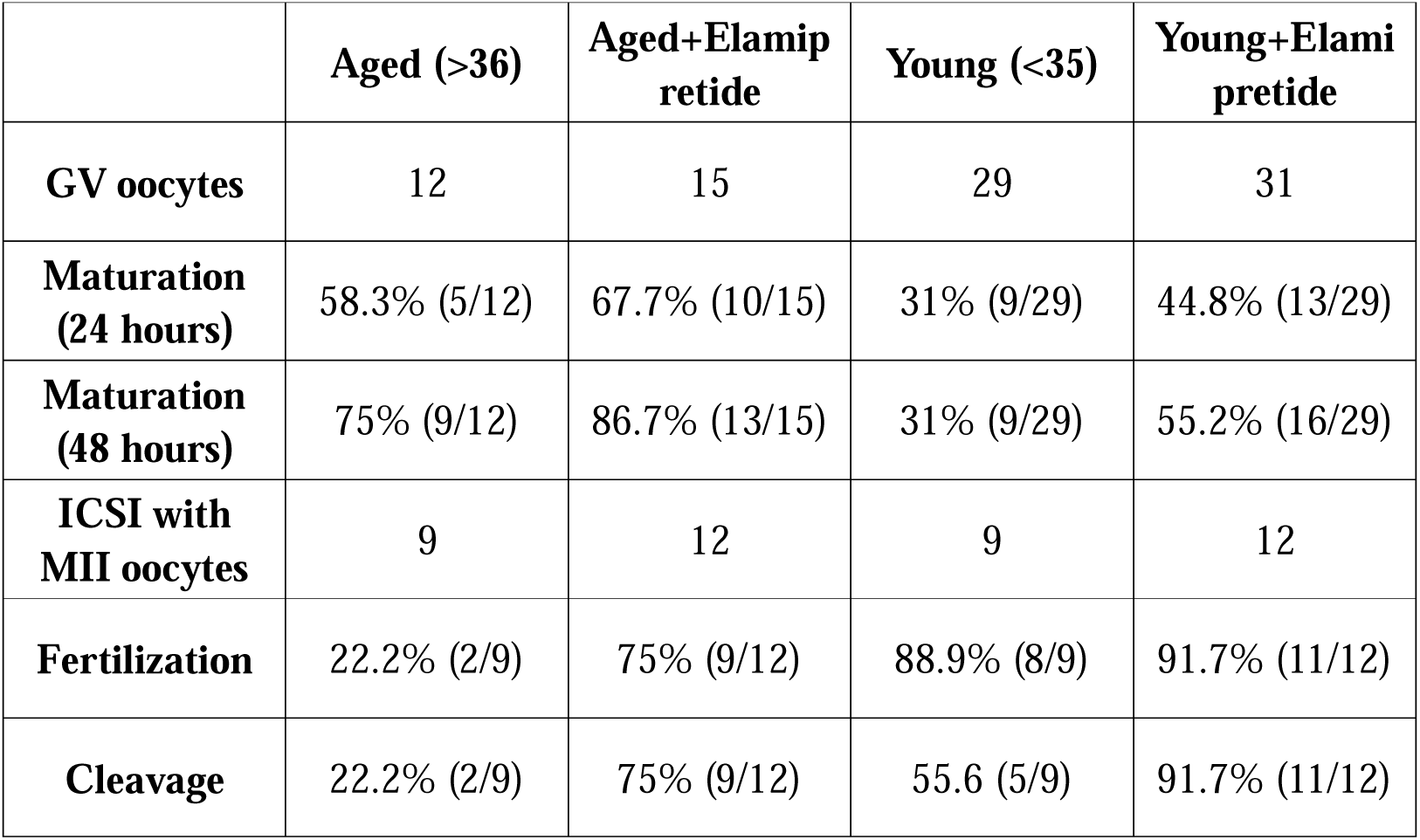
Rates of human oocyte maturation, fertilization and cleavage.

|  | **Aged (>36)** | **Aged+Elamip**<br>**retide** | **Young (<35)** | **Young+Elami**<br>**pretide** |
| --- | --- | --- | --- | --- |
| **GV oocytes** | 12 | 15 | 29 | 31 |
| **Maturation**<br>**(24 hours)** | 58.3% (5/12) | 67.7% (10/15) | 31% (9/29) | 44.8% (13/29) |
| **Maturation**<br>**(48 hours)** | 75% (9/12) | 86.7% (13/15) | 31% (9/29) | 55.2% (16/29) |
| **ICSI with**<br>**MII oocytes** | 9 | 12 | 9 | 12 |
| **Fertilization** | 22.2% (2/9) | 75% (9/12) | 88.9% (8/9) | 91.7% (11/12) |
| **Cleavage** | 22.2% (2/9) | 75% (9/12) | 55.6 (5/9) | 91.7% (11/12) |

### Short-term elamipretide treatment improves fertility of aged mice

An optimal therapeutic protocol for elamipretide in aged mice was initially established, with body weight and ovarian status systematically documented in both young and aged mice (Fig. S1A). As shown in Figure 1A, daily intraperitoneal elamipretide was administered to aged mice for 7 or 10 days, with age-matched and young controls receiving equivalent saline. Upon completion, mice were either naturally mated for fertility evaluation or super-ovulated with PMSG/hCG to harvest oocytes for subsequent assays. We assessed daily doses of 3, 5, and 10 mg/kg in 10-month-old aged mice and collected MII oocytes from the ampulla. The 5 mg/kg regimen produced the most optimal efficacy in increasing the number of ovulations (Fig. 1B). In Figure 1C, using 5 mg/kg/day, we treated 8-, 10-, and 12-month-old mice for 7 or 10 days and then assessed fertility. The 7-day regimen improved reproductive performance in all age groups, with the most pronounced gain observed in 10-month-old mice. And the beneficial effect of elamipretide on fertility was limited to a short-term window of only one month (Fig. S1B). Consequently, the subsequent studies were therefore performed in 10-month-old mice subjected to the 7-day elamipretide protocol. Ovarian sections likewise showed that elamipretide restored the number of dominant follicles and rejuvenated the aged ovary (Fig. 1D). After parthenogenetic activation of isolated MII oocytes, elamipretide markedly elevated the pronuclear formation rate in aged oocytes, confirming a substantial recovery of their fertilization potential (Fig. 1E). This was also consistent with its role on repairing cortical granule trafficking (Fig. S1C). Following fertilization, aged oocytes exhibited a reduced 2-cell rate and a markedly lower 4-cell rate than young controls, while elamipretide effectively reversed 2- to 4-cell progression, restoring early embryonic development (Fig. 1F). Therefore, elamipretide restores the dominant-follicle reserve in aged mice, increasing ovulation numbers while enhancing oocytes fertilization ability and rescuing the 2- to 4-cell developmental block, thereby collectively improving reproductive capacity in aging mice.

**Figure 1.**
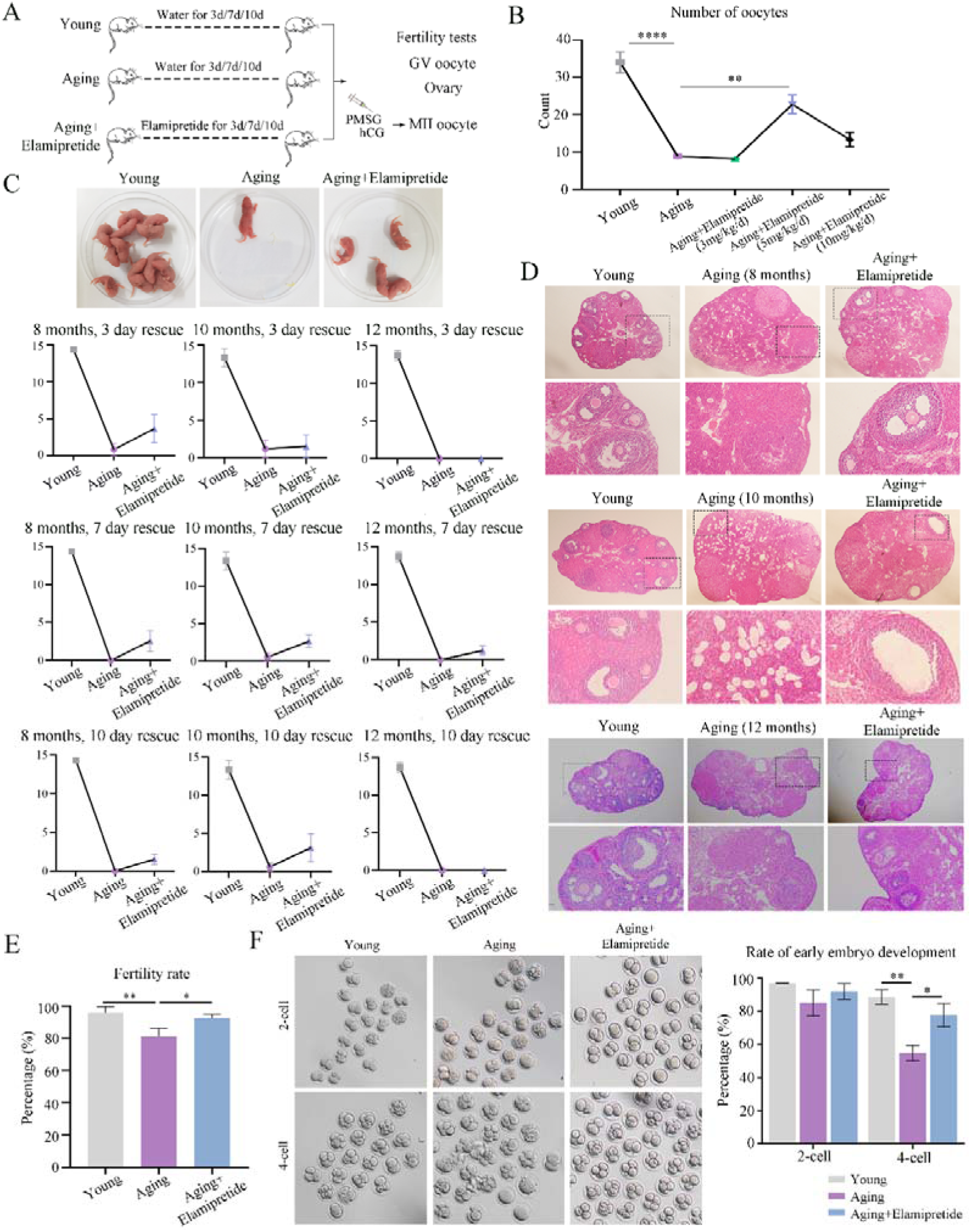
Short-term elamipretide treatment boosts fertility in aged mice. (A) Treatment regimen of elamipretide in aged mice. (B) Dose-escalation study to determine the optimal elamipretide regimen. (C) Fertility assessment across varying treatment durations. (D) Ovarian sections from 8-, 10- and 12-month-old mice after 7-day elamipretide treatment showed a marked increase in high-quality of follicles. (E) elamipretide restored pronucleus formation rates after parthenogenetic activation. (F) elamipretide effectively rescued the 2- to 4-cell transition. * P < 0.05, ** P < 0.01, **** P < 0.0001.

### Elamipretide reshapes ovarian Vitamin B6 metabolism in aged mice

We performed metabolomics to examine the metabolism change after elamipretide treatment. First, all detected metabolites were clustered based on their temporal dynamics, yielding 12 distinct expression patterns (Fig. 2A). In Figure 2B, compared with the young group, aged ovaries exhibited 250 differentially accumulated metabolites (122 up, 128 down), whereas elamipretide treatment shifted 30 metabolites relative to aged group (12 up, 18 down). And we performed principal-component analysis on the complete metabolome and the differential metabolite subset, uncovering their overall distribution patterns and key discriminating dimensions (Fig. 2B). The heat map revealed an age-dependent metabolite signature that elamipretide restored to a young-like profile, and the Venn diagram revealed 11 differential metabolites shared between the two comparison groups (Fig. 2C). Figure 2D illustrated that metabolite abundance was markedly reduced in aged mice yet effectively restored to youthful abundance following elamipretide treatment. We performed GO-based enrichment clustering on the filtered metabolome and plotted the top 14 most over-represented metabolic categories as a bar chart (Fig. 2E). We then generated correlation networks for the differential metabolites identified in young vs. aging and aging vs. aging+elamipretide comparisons to visualize their interrelationships and regulatory hubs (Fig. 2F). We further mapped the differential metabolites to KEGG pathways, extracting the top 10 enriched routes for each pairwise comparison, with the vitamin B6 pathway showing a pronounced alteration (Fig. 2G). Additionally, we constructed a heat map showing the relative abundance of the 11 shared metabolites across the different groups (Fig. 2H). We next selected the most dramatically altered metabolites, LAC (L-Acetylcarnitine) and AVA (Aminovaleric acid). Individual or in combined treatment with them failed to restore first polar body extrusion in aged oocytes, whereas VB6 supplementation fully reversed the age-associated decline (Fig. 2I). Therefore, elamipretide thus appears to enhance the maturational quality of aged oocytes by modulating vitamin B6 metabolism.

**Figure 2.**
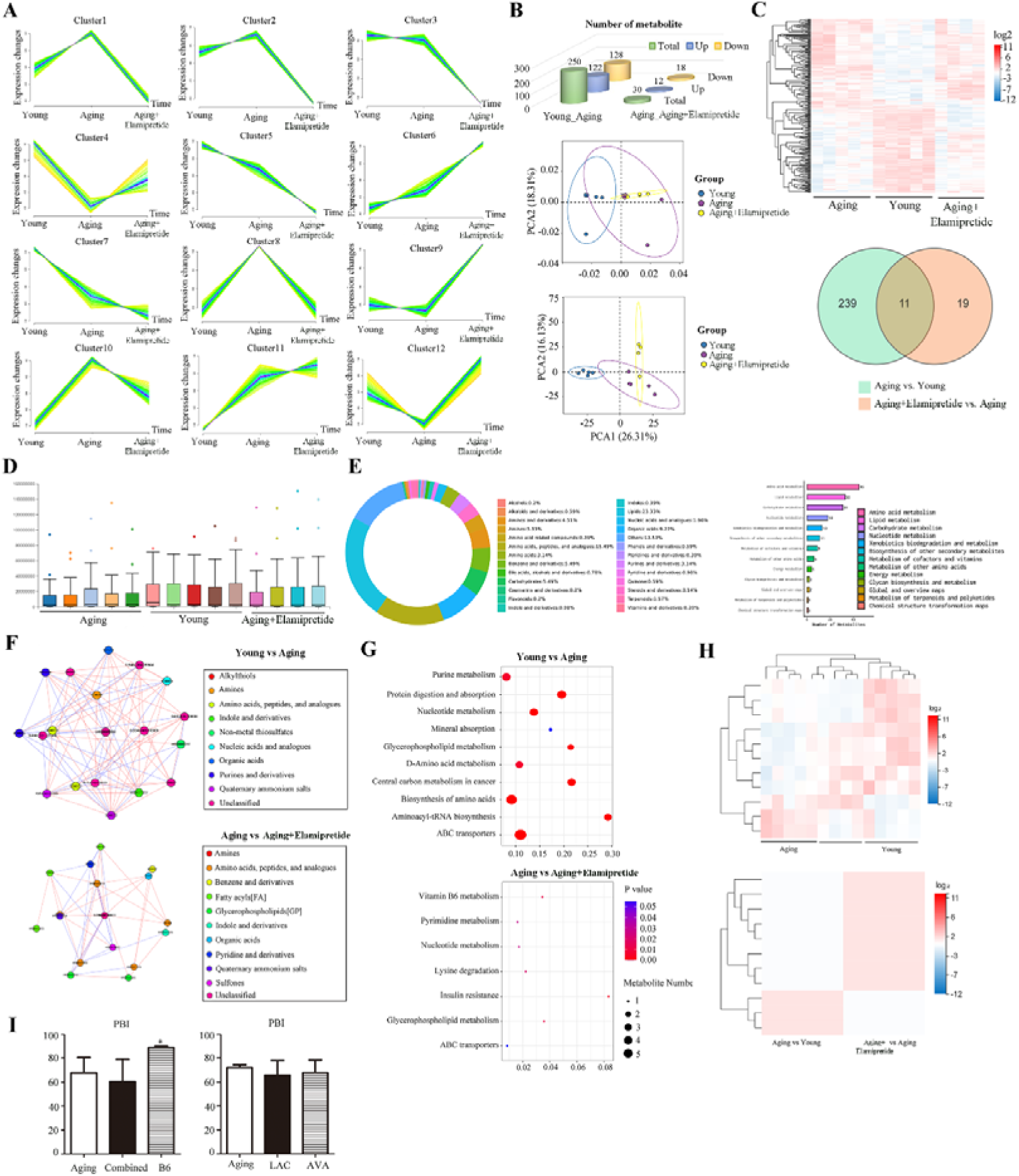
Elamipretide reshapes ovarian Vitamin B6 metabolism in aged mice. (A) Metabolite expression profiles across sample groups were clustered, grouping metabolites with similar expression patterns into the same cluster. (B) The number of up- and down- regulated differential metabolites were tailed, and PCA dimensionality reduction was used to visualize the inter-group separation trend. (C) Hierarchial clustering of differential metabolites directly revealed their expression patterns across the three sample groups, 11 differential metabolites were shared between the two comparison groups. (D) Global abundance profiling of all detected metabolites across every group. (E) Circular chart of metabolite categories and bar chart of metabolic pathway classification. (F) Correlation network map of differential metabolites. (G) Bubble plot of metabolic pathway enrichment analysis. (H) Hierarchical clustering of the 11 shared differential metabolites. (I) LAC, AVA, and vitamin B6 rejuvenated aged oocyte developmental competence. * P < 0.05.

### Elamipretide alters general transcript levels in mouse aged oocytes

We collected the oocytes for transcriptome analysis, with 15 oocytes each group. In the aging group compared with the young group, there was a total of 829 genes changed, with 420 upregulated and 409 genes downregulated. While in the elamipretide treatment group compared with the aging group, of the 2,081 differentially expressed genes, 965 were upregulated and 1,116 were downregulated (Fig. 3A). In the Figure 3B, it was showed that between these two sets of differentially expressed genes, 518 genes were commonly shared. The volcano plot displays the fold changes of differentially expressed genes and distinguishes the upregulated from the downregulated. The heatmap clusters differentially expressed genes, allowing us to visualize the overall expression patterns across the young, the aging and the aging+elamipretide groups (Fig. 3C). Then we performed GO enrichment analysis on the significantly differentially expressed genes in Figure 3D. We found that elamipretide treatment alleviates alterations in membrane-bound organelles and lipid transport and metabolism processes. Subsequent KEGG analysis of the differentially expressed genes reveals that the VEGF signaling pathway is markedly altered in oocytes from aged mice compared with the young, and the VEGF pathway expression is reversed in elamipretide treated mouse oocytes (Fig 3E). Therefore, elamipretide may modulate the maturation quality of aged oocytes via the VEGF pathway.

**Figure 3.**
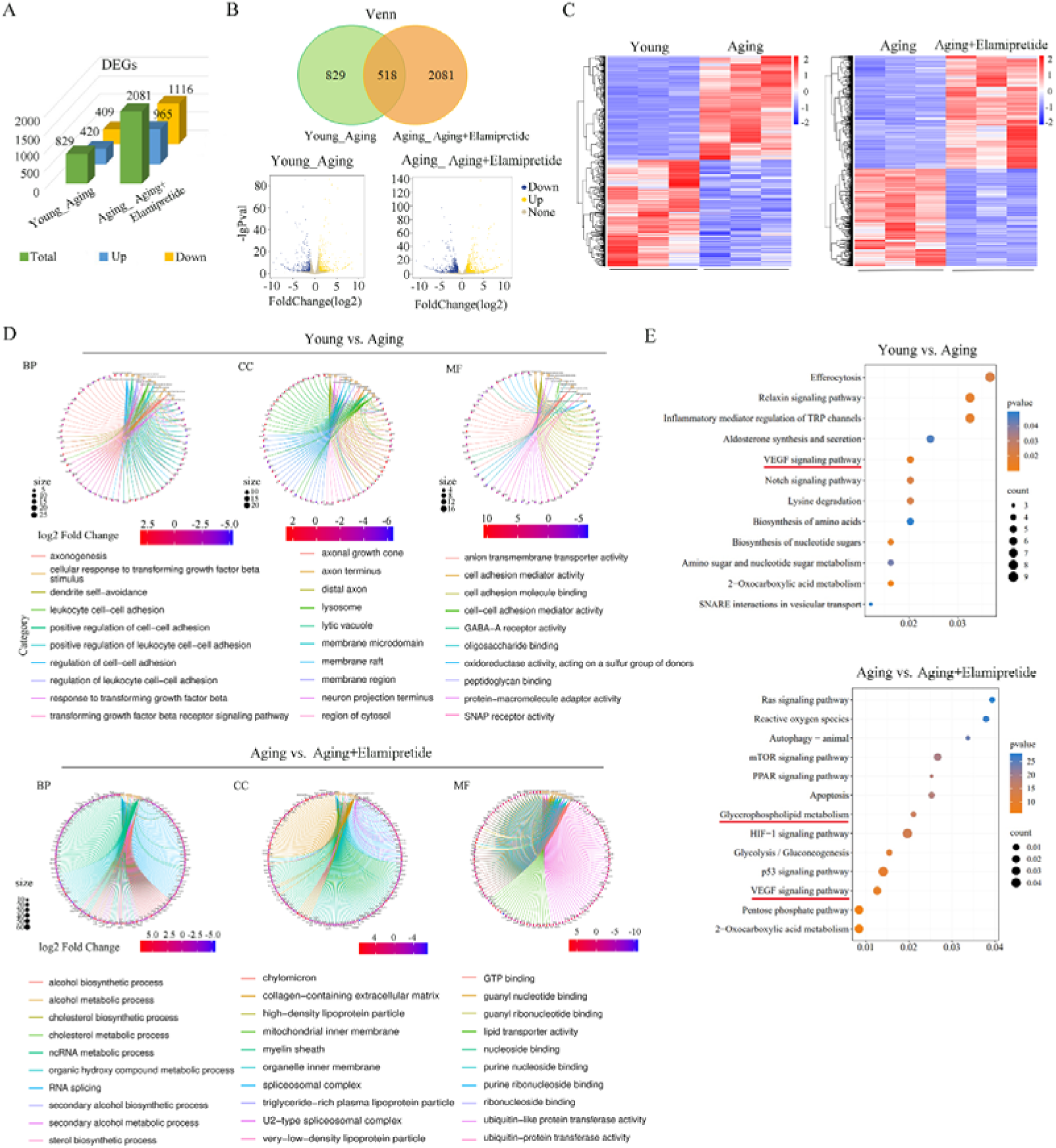
Elamipretide alters transcript levels in mouse oocytes. (A) Differential gene expression showed bidirectional regulation, with up- and down- regulated genes clearly segregated. (B) Venn and volcano plots of differentially expressed genes revealed 518 shared genes. (C) Heat map of differentially expressed genes. (D) Chord diagram of GO enrichment clusters for differentially expressed genes. (E) Top 12 KEGG pathways enriched by differentially expressed genes shown in a bubble plot.

### Elamipretide promotes cytoskeletal dynamics and spindle migration of aged oocytes

Using GO analysis, we identified cytoskeleton-related biological processes rescued by elamipretide verses aged oocytes (Fig. 4A). In the Figure 4B, we first examined α-tubulin assembly and chromosome alignment by immunofluorescence, which showed that aged oocytes displayed aberrant spindles and scattered chromosomes, whereas elamipretide reduced the frequency of these anomalies. γ-tubulin was used to mark the spindle poles. Our staining revealed that aged oocytes lost γ-tubulin foci at the poles, and elamipretide restored them (Fig. 4C). Acetylated tubulin is commonly used as a marker of microtubule stability. Our immunofluorescence staining results showed that aging decreased the level of tubulin acetylation, indicating that aging compromises microtubule stability, whereas supplementation with elamipretide restored tubulin acetylation to normal levels (Fig. 4D). Additionally, we assessed whether actin-network organization was compromised. Cortical actin assembly remained intact in aged oocytes, and elamipretide treatment did no significant alteration (Fig. 4E). Notably, cytoplasmic actin-network assembly was disrupted by aging, and this decline was effectively reversed by elamipretide supplementation (Fig. 4F). Subsequently, we detected the expression levels of related proteins via western blot analysis. The results demonstrated that Arp2, a protein closely associated with cortical actin assembly, was significantly downregulated in aged oocytes, and treatment with elamipretide failed to reverse the decreased expression of this protein (Fig. 4G). ROCK1 is involved in regulating the assembly of cytoplasmic actin in oocytes, and our western blot results confirmed that elamipretide can effectively reverse the downregulation of ROCK1 protein expression induced by aging (Fig. 4H). Furthermore, protein analysis further confirmed that SS□31 effectively rescued the aging□induced decrease in microtubule acetylation, and the expression level of acetylated tubulin (Ac□tubulin) was significantly restored in aged oocytes following SS□31 treatment (Fig. 4H). Thus, elamipretide sustains spindle assembly and migration in aged oocytes by elevating microtubule acetylation and promoting cytoplasmic actin nucleation.

**Figure 4.**
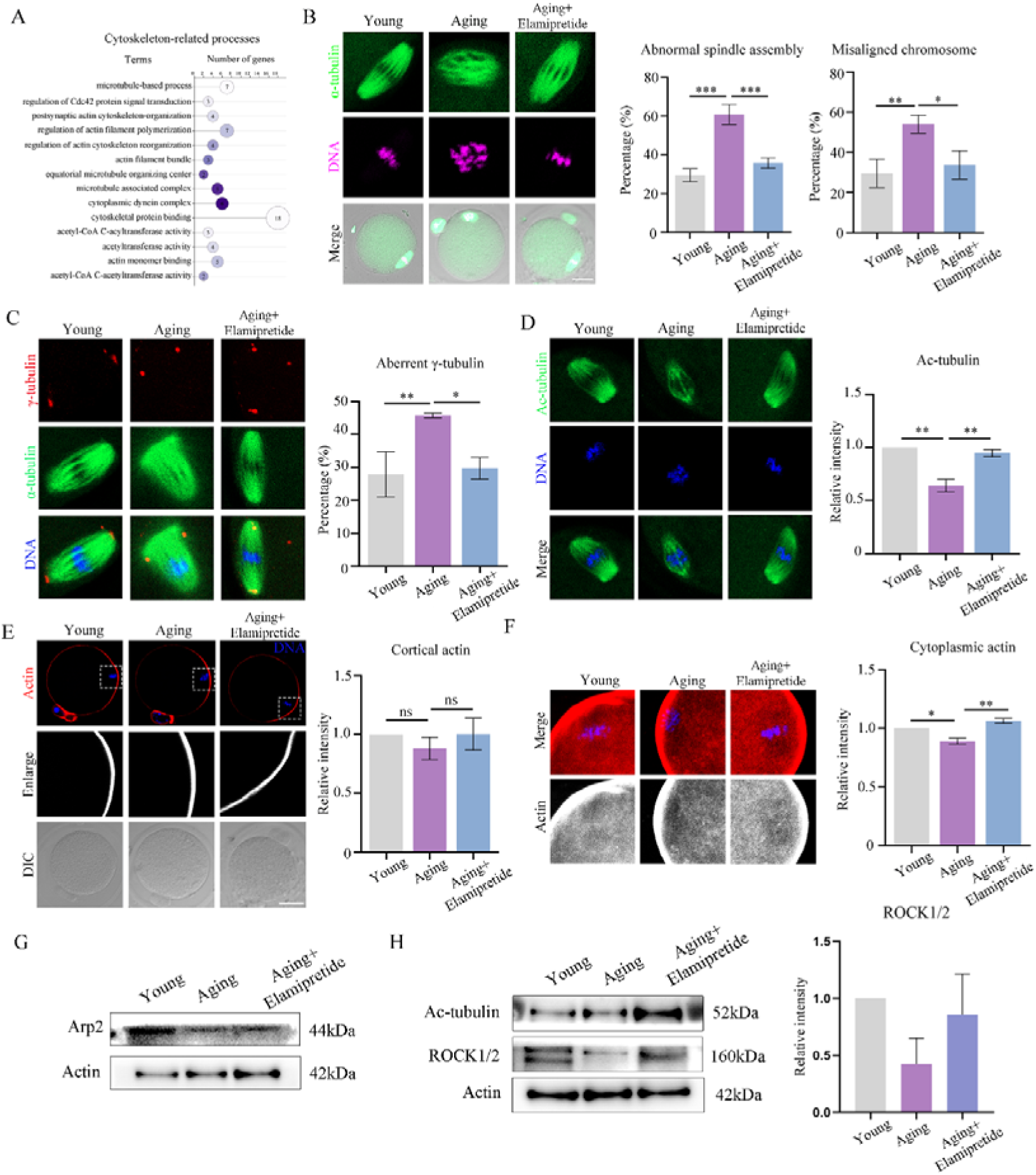
Elamipretide promotes cytoskeletal assembly and spindle migration. (A) GO classification of cytoskeleton-related processes improved by elamipretide treatment in aged oocytes. (B) elamipretide restored orderly spindle assembly and corrects chromosome alignment. Green, α-tubulin; purple, DNA; scale bar, 20 μm. (C) elamipretide re-established the proper localization of γ-tubulin. Red, γ-tubulin; green, α-tubulin; blue, DNA; scale bar, 20 μm. (D) elamipretide boosted acetylated tubulin levels, stabilizing the meiotic spindle in aged oocytes. Green, α-tubulin; blue, DNA; scale bar, 20μm. (E) Cortical F-actin remained unchanged in aged oocytes. (F) The cytoplasmic actin was significantly reduced in aged oocytes, and elamipretide fully restored cytoplasmic actin levels. Red, actin; blue, DNA; scale bar, 20μm. * P < 0.05, ** P < 0.01, *** P < 0.001, and ns, no significant difference.

### Elamipretide alleviates mitochondria-driven oxidative stress in aged oocytes

Oocyte maturation extends beyond nuclear progression to include cytoplasmic maturation, with mitochondrial rearrangement as its pivotal event. From the transcriptome analysis, we first extracted the nine most significantly enriched GO pathways distinguishing the aging group from the aging+elamipretide group. In Figure 5A, differentially expressed genes were chiefly enriched for inner mitochondrial membrane organization, mitochondrial metabolism, and oxidative-stress response. We selected a panel of mitochondria-related key genes and quantified the mRNA levels by qPCR, elamipretide fully reversed the age-associated transcriptional changes (Fig. 5B). In Figure 5C, the fluorescence imaging of MII oocytes revealed that aging displaced mitochondria from the peri-spindle domain and drove their condensation into cytoplasmic clusters, while elamipretide restored mitochondrial positioning. Mitochondrial copy number and ATP synthesis level reflected the organelle’s quantity reserve and energetic capacity, respectively. We found that elamipretide normalized the supraphysiological ATP synthesis to young levels and partially reinstated mitochondrial DNA copy number (Fig. 5D). TMRE is a fluorescent probe for quantifying mitochondrial membrane potential. We found that mitochondrial membrane potential declined in aged oocytes, likely accounting for the reduced ATP production, and elamipretide treatment restored potential to a higher level (Fig. 5E). Therefore, we examined the expression levels of mitochondrial fission proteins INF2 and DRP1. The results showed that INF2 expression was significantly decreased in aging oocytes, and SS□31 treatment failed to restore its expression level (Figure 5F). In contrast, Western blot results and statistical analysis confirmed that SS□31 significantly upregulated DRP1 expression in aging oocytes (Fig. 5G). Mitochondrial depolarization commonly signals intensified oxidative stress and ROS burst, so we next quantified the oxidative-stress burden in oocytes. We profiled a panel of oxidative-stress-related genes and found that elamipretide partially reversed their age-induced overexpression (Fig. 5H). Fluorescence imaging revealed that elamipretide markedly restored redox homeostasis in aged oocytes (Fig. 5I). Additionally, we examined oxidative-stress-induced apoptosis in aged oocytes. elamipretide only partially reversed the transcriptional changes of apoptosis-related genes (Fig. 5J), and fluorescence staining confirmed that it failed to rescue the elevated apoptosis caused by aging (Fig. 5K). The findings indicate that elamipretide can restore oxidative stress by improving mitochondrial distribution and function in aged oocytes.

**Figure 5.**
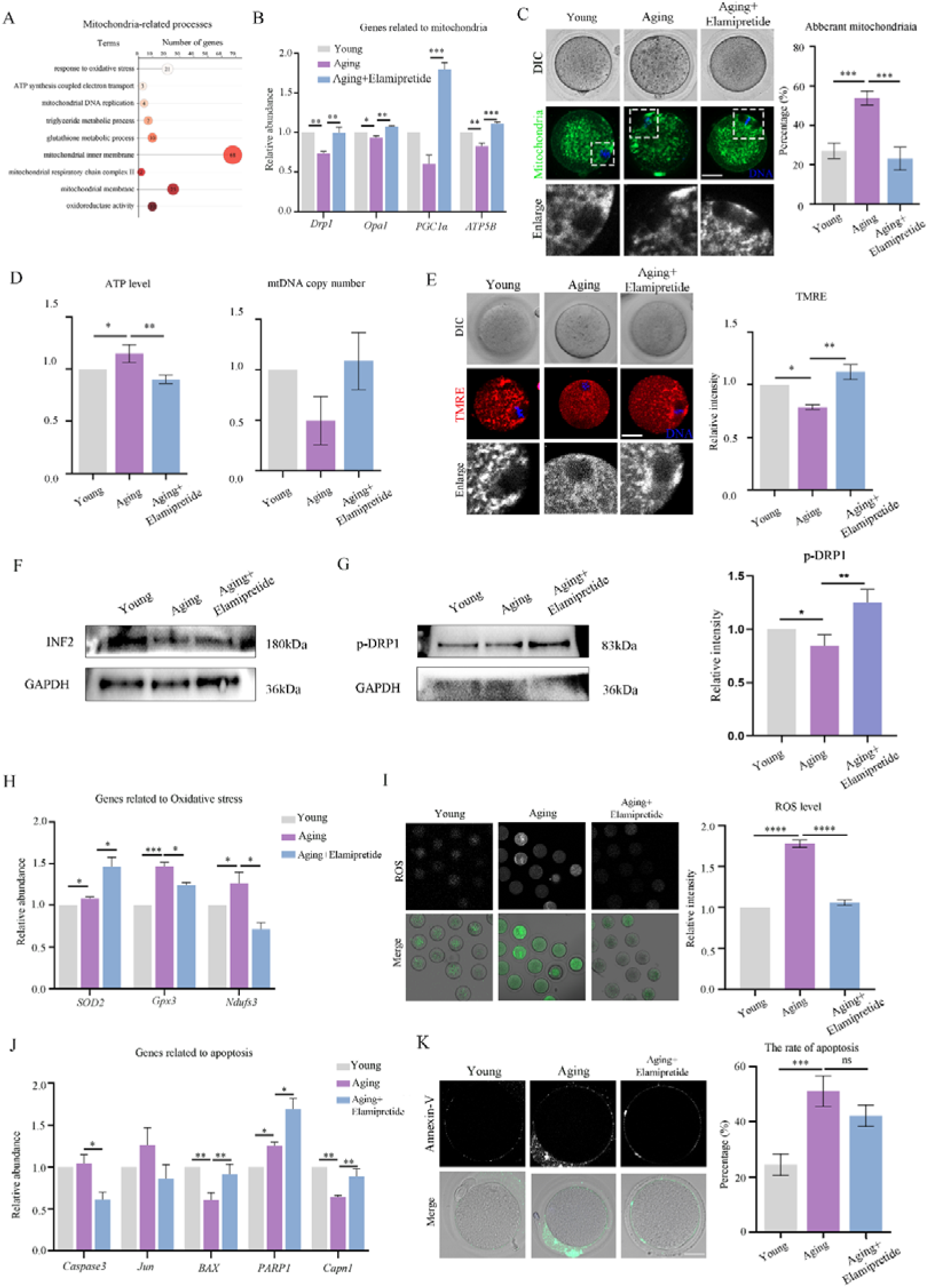
Elamipretide alleviates mitochondria-driven oxidative stress. (A) GO clusters related to mitochondrial metabolism identified between Aging + elamipretideversus aged oocytes. (B) Quantification of mRNA levels for mitochondria-related functional genes. (C) elamipretide corrected the aberrant peri-spindle distribution of mitochondria in aged oocytes. Green, mitochondria; blue, DNA; scale bar, 20μm. (D) elamipretide restored overly elevated ATP to physiological levels and simultaneously boosted mitochondrial number. (E) TMRE reported mitochondrial function, and elamipretide ameliorated mitochondrial dysfunction by elevating mitochondrial membrane potential. Red, TMRE; blue, DNA; scale bar, 20 μm. (F) Changes in mRNA expression of oxidative-stress-related genes. (G) elamipretide brings excessive oxidative stress in aged oocytes back to youthful levels. Green, ROS; scale bar, 20μm. (H) Alterations in mRNA levels of apoptosis-related genes. (I) elamipretide failed to rescue oxidative-stress-induced apoptosis in aged oocytes. Green, Annexin-V; scale bar, 20μm. * P < 0.05, ** P < 0.01, *** P < 0.001, **** P < 0.0001, and ns, no significant difference.

### Elamipretide restores organelle-associated processes in aged oocytes

Cytoplasmic maturation also encompasses the remodeling and functional maturation of organelles such as the endoplasmic reticulum and Golgi apparatus. By screening membrane-organelle pathways, we found that ER-, Golgi- and lysosome-related processes were all perturbed (Fig. 6A). To assess the impact of elamipretide on Golgi function, we mapped Rab10 localization in oocytes by immunofluorescence. We found that elamipretide fully reversed the age-dependent decline in Rab10 expression, restoring it to normal levels (Fig. 6A). In Figure 6B, analysis of Golgi-related transcripts revealed that elamipretide rescued the age-dependent decline in the functional marker GM130, restoring its expression to normal levels. Our protein detection results further confirmed that SS□31 effectively restored the expression levels of GM130 and Rab10 in aged oocytes, thereby improving Golgi function in aged oocytes (Fig. 6C). The ER also plays a pivotal role in oocyte cytoplasmic maturation. With aging, ER strands abandoned their spindle-adjacent stronghold and scattered evenly throughout the cytoplasm. And elamipretide returned them to the subcortical spindle zone (Fig. 6D). Further analysis of the ER stress marker GRP78 revealed that elamipretide effectively reversed its age-dependent decline, indicating that the peptide rescued insufficient ER stress adaptation (Fig. 6E). Our western blot analysis confirmed that SS□31 effectively restored the downregulated expression of GRP78 in aged oocytes, thereby improving the endoplasmic reticulum stress response in oocytes (Fig. 6F). Following elamipretide intervention, transcript levels of the ER-related genes GRP78, CHOP and ATF4 returned to normal, further confirming the peptide’s restorative effect on ER function (Fig. 6G). Additionally, lysosome-mediated autophagic activity constitutes an equally indispensable facet of cytoplasmic maturation. In aged oocytes autophagic vacuoles accumulated abnormally and lysosomes lost their normal distribution, and elamipretide restored the even cytoplasmic pattern (Fig. 6H). In senescent oocytes LC3-positive vesicles were fewer and scattered, elamipretide treatment amplified LC3 fluorescence and reinstated autophagic activity (Fig. 6I). The statistical analysis data proved the results in Figure 6J. The western blot analysis demonstrated that the expression level of LC3 was slightly decreased in aged oocytes, and supplementation with SS□31 restored its expression to the level of young oocytes (Fig. 6K). Transcript profiling of autophagy genes showed elamipretide returned their mRNA abundance to the normal range (Fig. 6L). The results demonstrate that elamipretide orchestrates organelle repositioning and concomitantly restores protein synthesis and autophagic flux in aged oocytes.

**Figure 6.**
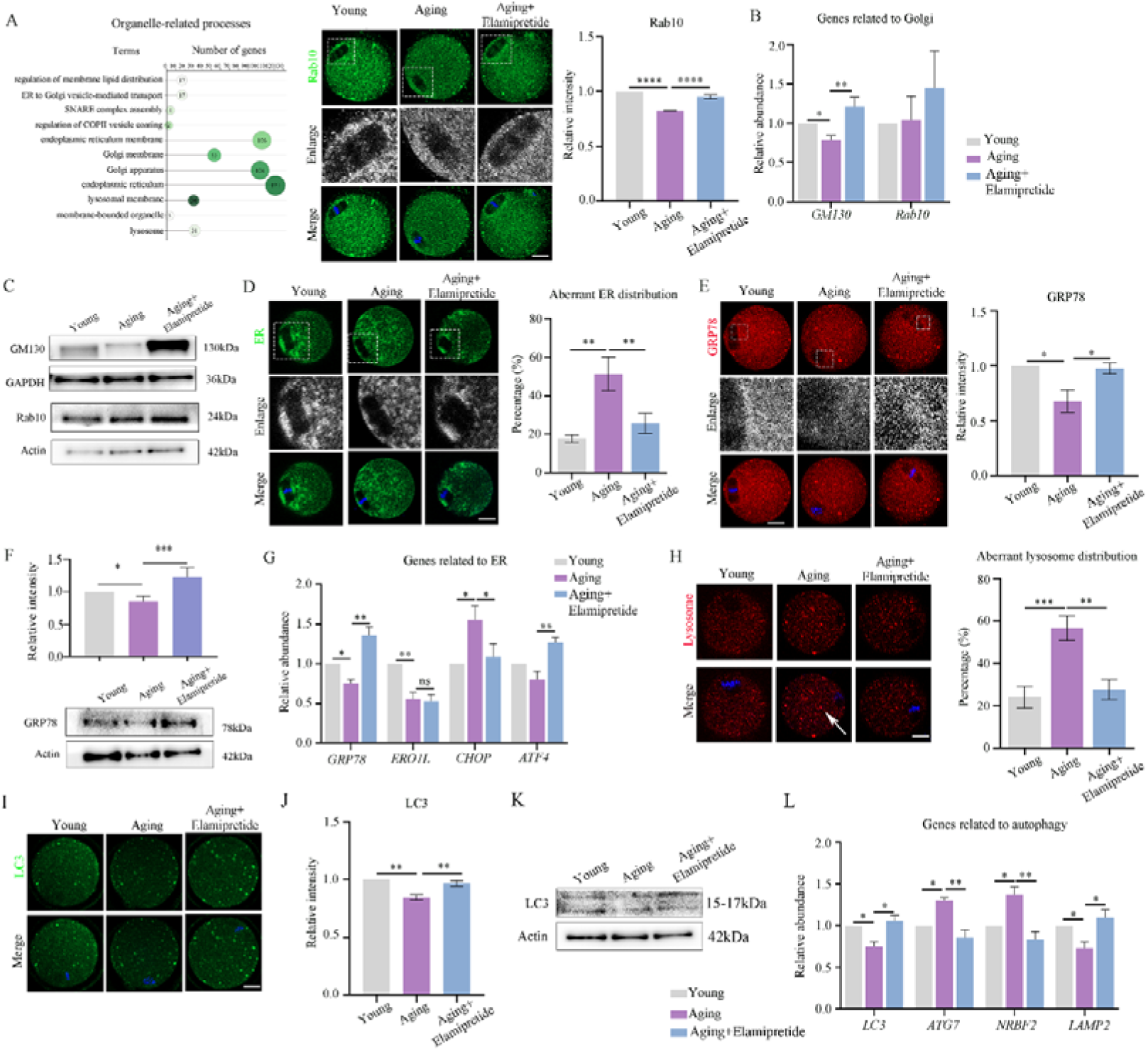
Elamipretide restores organelle-associated processes. (A) GO analysis of organelle-related processes and fluorescence intensity analysis of the Golgi functional protein Rab10after elamipretide treatment. Green, Rab10; blue, DNA; scale bar, 20μm. (B) Detection of mRNA expression levels of Golgi-associated genes. (C) elamipretide restored the loss of ER localization around the spindle in aged oocytes. Green, ER; blue, DNA; scale bar, 20μm. (D) The ER stress protein GRP78 was aberrantly reduced in aged oocytes, and elamipretide restored it to normal levels. Red, GRP78; blue, DNA; scale bar, 20μm. (E) Changes in the mRNA expression levels of ER-related genes. (F) elamipretide alleviated the excessive accumulation of lysosomal autophagic vesicles in oocytes caused by aging. Red, lysosome; blue, DNA; scale bar, 20μm. (G) elamipretide recovered the abnormal reduced LC3 expression levels in aged oocytes. Green, LC3; blue, DNA; scale bar, 20μm. (H) The mRNA expression levels of autophagy-related genes were significantly improved after elamipretide treatment. * P < 0.05, ** P < 0.01, *** P < 0.001, **** P < 0.0001, and ns, no significant difference.

### Elamipretide restores aged oocyte maturation competence with IVM system in mice

Considering the actual situation of IVF (*in vitro* fertilization) center for aged oocyte culture, we concurrently evaluated whether elamipretide can improve the maturation defects of aged oocytes under *in vitro* culture conditions. We established three concentration gradients of 100, 200 and 400 μM to determine the optimal rescue concentration. It is showed that exogenous elamipretide repaired the developmental defects of oocytes caused by aging (Fig. 7A), with 200 μM being the optimal rescue concentration (Fig. 7B). Subsequent phenotypic analyses were therefore conducted at this concentration. We further found that when MII-stage oocytes were collected and subjected to in vitro fertilization, supplementation of elamipretide during culture significantly improved subsequent embryonic development, as evidenced by an increased blastocyst rate (Fig. S2A). Statistical analysis confirmed this effect (Fig. S2B). Cortical-granule (CG) translocation is a cardinal indicator of cytoplasmic maturation and subsequent developmental potential. Here we showed that aging severely disrupted CG trafficking, producing aberrant peri-cortical clustering, and elamipretide supplementation fully reinstated the monolayer beneath the cortex (Fig. S2C). The data analysis also underscoring the capacity of elamipretide to rectify the vesicular trafficking defects inherent to aged oocytes (Fig. S2D). To delineate the age-evoked transcriptional landscape, we harvested in-vitro-matured oocytes synchronously arrested at metaphase I and subjected them to transcriptome profiling. A total of 1,255 genes exhibited significant differential expression between young and aging groups; upon elamipretide supplementation, 317 genes remained differentially expressed relative to the aging group (Fig. S3A). Of these, 114 differentially expressed genes were common to both comparisons, and the heatmap of differential genes between the two comparison groups was presented in Figure S3B. Subsequent KEGG and Gene Ontology analyses revealed that these differentially expressed genes were significantly enriched in pathways related to organelle biogenesis and mitochondrial metabolism (Fig. S3C). GO enrichment clustered the DEGs into functional modules, revealing that elamipretide improved oocyte defects in energy metabolism, apoptosis regulation, and protein synthesis (Fig. S3D). We identified the top 10 significantly differentially expressed cytoskeleton-related genes from the transcriptome data (Fig.7C). Fluorescence staining revealed that exogenous elamipretide reversed age-induced chromosome misalignment and spindle migration failure (Fig. 7D). Besides, elamipretide largely corrected the cytoskeletal disorganization caused by aging, which brought microtubule acetylation back to normal and stabilized the microtubule in aged oocytes cultured in vitro (Fig. S4A). The aberrant γ-tubulin distribution observed was likewise ameliorated by elamipretide supplementation (Fig. S4B), and statistical analysis confirmed this effect (Fig. S4C). However, elamipretide failed to restore the reduced cortical F-actin density of aged oocytes to the level seen in controls (Fig. S4D). We additionally identified differentially expressed genes associated with the mitochondrial-apoptotic pathway (Fig. 7E). In Figure 7F, aged oocytes exhibited aberrantly high mitochondrial fluorescence after in vitro culture, which was markedly attenuated by elamipretide treatment. In addition, elamipretide effectively attenuated the elevated oxidative stress characteristic of aged oocytes (Fig. 7G). elamipretide likewise normalized the age-induced hyperpolarization-driven surge in TMRE fluorescence, returning mitochondrial membrane potential to levels compared to those of the control group (Fig. S5A). And elamipretide curbed the Parkin-dependent mitophagy surge seen in aged IVM oocytes (Fig. S5B). We found that elamipretide supplementation during IVM alleviated oxidative stress in aged oocytes (Fig. S5C), yet failed to normalize the transcription of related genes (Fig. S5D). Moreover, the elevated apoptotic signaling induced by aging was effectively ameliorated by elamipretide supplementation during IVM (Fig. S5E), the transcription levels of the relevant genes were also restored (Fig. S5F). The genes related to organelles shown in Figure 7H were selected from the transcriptome data. We mapped Golgi localization in oocytes and observed that elamipretide reinstated its peri-spindle enrichment, a positioning abolished by aging (Fig. 7I). The statistical analysis confirmed this finding (Fig. S6A). Examination of the Golgi marker Rab10 revealed that elamipretide boosted its spindle-proximal accumulation, signifying functional recovery of the Golgi apparatus (Fig. S6B). The ER formed large aggregated masses in aged oocytes, and elamipretide treatment restored its proper localization (Fig. 7J), which was proved by the statistical data (Fig. S6C). And elamipretide reduced elevated GRP78 in aged oocytes, which relieved ER stress and restored ER function (Fig. S6D). Additionally, elamipretide reversed the dispersed distribution of autolysosomes that were aberrantly clustered in age oocytes during IVM (Fig. S6E). LC3 puncta intensity was also visibly reduced, indicating diminished autophagosome formation (Fig. S6F). Our findings demonstrated that elamipretide effectively rescued the cytoplasmic-maturation failure characteristic of aged oocytes during *in vitro* culture, restoring organellar reorganization, cytoskeletal architecture and developmental competence to levels approaching those of the young group.

**Figure 7.**
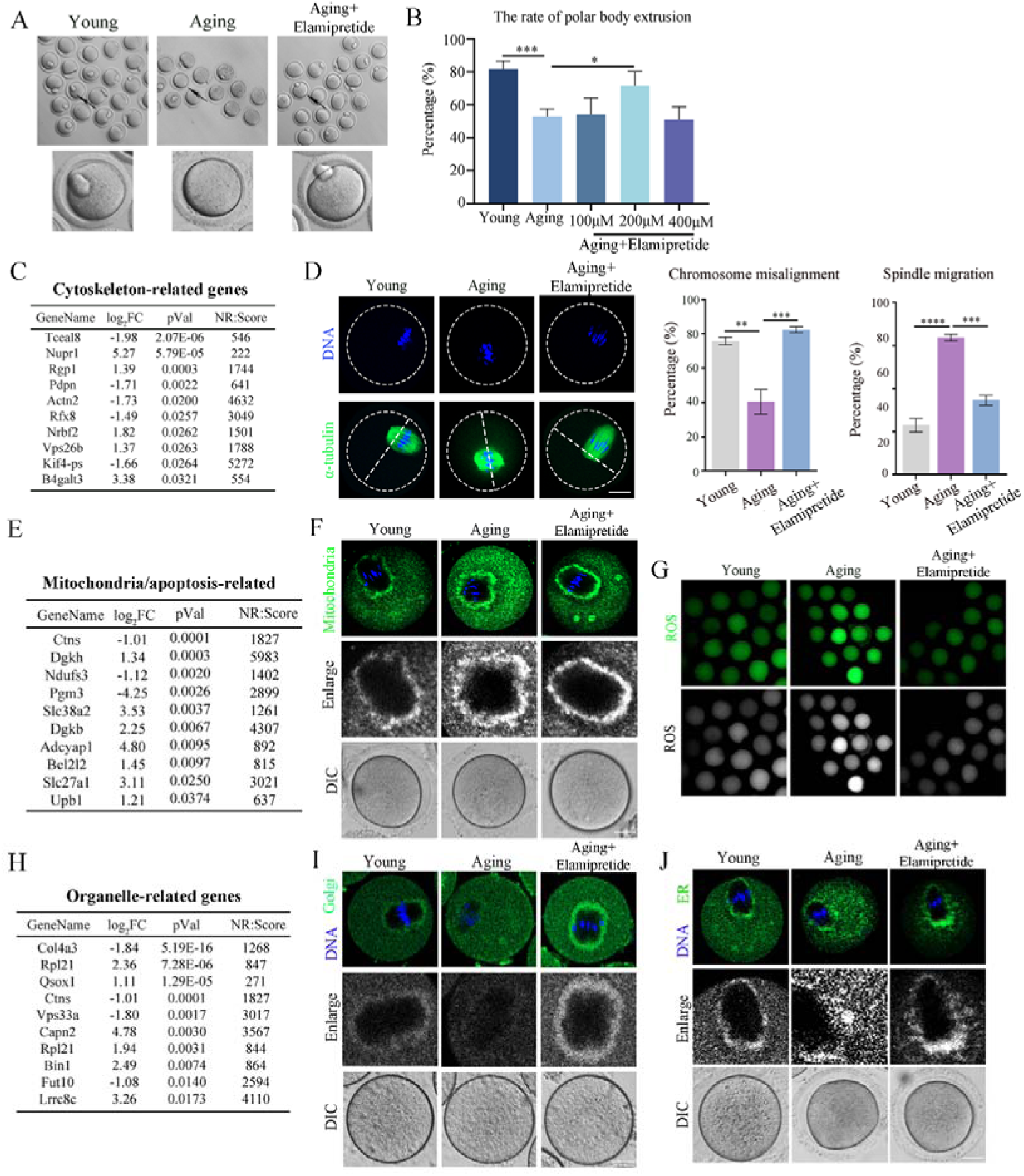
IVM with elamipretide restores maturation competence in aged oocytes. (A) Supplementation of elamipretide during in vitro culture improved oocyte maturation quality. (B) elamipretide rescued oocytes in a concentration-gradient assay, with 200μM identified as the optimal dose. (C) Cytoskeleton-related genes showed significant differential expression after in vitro elamipretide supplementation. (D) In vitro supplementation of elamipretide improved chromosome alignment and spindle migration. Green, α-tubulin; blue, DNA; scale bar, 20μm. (E) Mitochondria-mediated apoptosis-related genes showed significant differential expression after in vitro elamipretide supplementation. (F) In vitro elamipretide supplementation alleviated aging-induced excessive accumulation of mitochondria around the spindle. Green, mitochondria; blue, DNA; scale bar, 20μm. (G) elamipretide alleviated the aging-induced elevation of ROS level. Green, ROS; scale bar, 20μm. (H) Organelle-related genes are significantly differentially expressed following in vitro elamipretide supplementation. (I) In vitro elamipretide supplementation restored the Golgi apparatus to its characteristic peripheral-spindle localization. Green, Golgi; blue, DNA; scale bar, 20μm. (J) In aged oocytes, the ER was aberrantly aggregated, and elamipretide supplementation restored its proper distribution. Green, ER; blue, DNA; scale bar, 20μm. * P < 0.05, ** P < 0.01, *** P < 0.001, **** P < 0.0001.

### Elamipretide improves post-ovulatory aged oocyte quality with IVM system in pigs

We then employed porcine oocytes to determine whether elamipretide likewise promoted maturation across species. As shown in Figure 8A, H_2_O_2_ exposure markedly reduced PBI extrusion in porcine oocytes, whereas elamipretide supplementation fully restored maturation to control levels. And 1mM emerged as the optimal treatment concentration (Fig. 8B). We found that elamipretide reduced the oxidative-stress level of H_2_O_2_-treated porcine oocytes (Fig. 8C), and the statistical analysis aligned with the staining results (Fig. 8D). Given the link between oxidative stress and mitochondrial function, we examined mitochondrial distribution in porcine oocytes and found that elamipretide reversed the H_2_O_2_-induced loss of mitochondria (Fig. 8E). Moreover, LC3 expression was restored after elamipretide treatment in H_2_O_2_-induced oocytes (Fig. 8F). elamipretide also repaired the abnormally high apoptosis induced by H_2_O_2_ treatment back to normal levels (Fig. 8G). Our results demonstrated that the ability of elamipretide to alleviate oxidative stress and restore oocyte developmental quality is functionally conserved across species.

**Figure 8.**
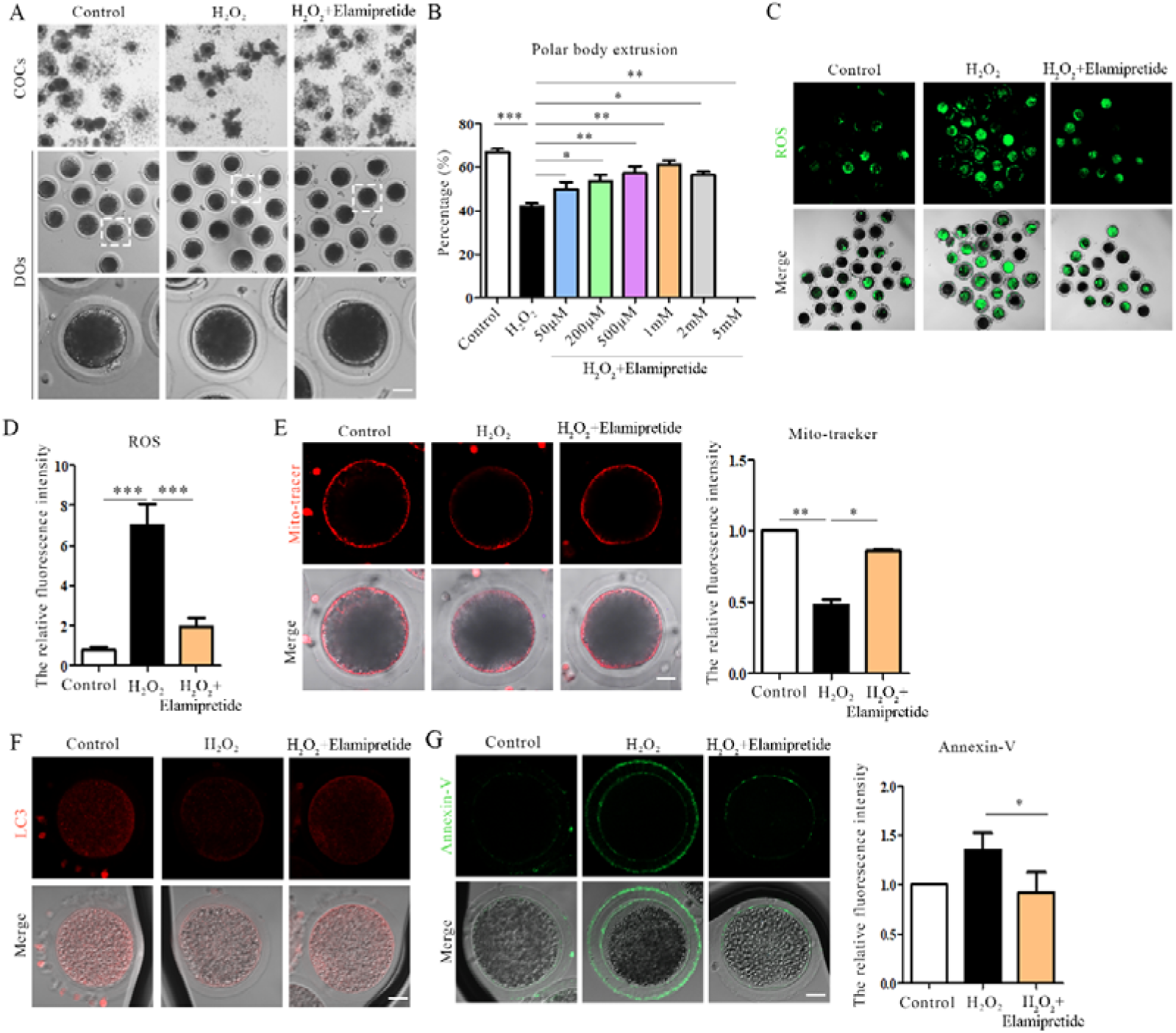
Elamipretide improves oocyte quality in pigs. (A) Elamipretide improved the maturation quality of porcine oocytes treated with hydrogen peroxide. (B) A concentration-gradient assay identified 1 mM as the optimal treatment concentration. (C) The abnormally elevated ROS induced by hydrogen peroxide treatment was significantly alleviated after elamipretide therapy. Green, ROS; scale bar, 20μm. (D) The statistical analysis of ROS level in porcine oocytes. (E) elamipretide reversed the mitochondrial fluorescence intensity reduced by hydrogen peroxide treatment to normal levels. Red, mitochondria; scale bar, 20μm. (F) elamipretide reversed hydrogen-peroxide-induced autophagy suppression. Red, LC3; scale bar, 20μm. (G) elamipretide mitigated the apoptosis surge triggered by oxidative stress in porcine oocytes. Green, Annexin-V; scale bar, 20μm. * P < 0.05, ** P < 0.01, *** P < 0.001.

### Elamipretide reverses aged oocyte quality via VEGF-dependent pathway

As shown in Figure 9A, transcriptome profiling identified VEGF-pathway genes with significant differential expression, and a heatmap was generated. We then validated these genes by qRT-PCR, confirming the expression trends observed in the transcriptome data (Fig. 9B). Western blot analysis further revealed that aging elevated VEGF-pathway protein levels, and elamipretide treatment effectively regained them to baseline (Fig. 9C). We first interrogated the PI3K/AKT axis with LY294002 and Wortmannin. In Figure S7A, 4μM LY294002 was selected as the working concentration, hyper-assembled cytoplasmic microfilaments were normalized, yet cortical F-actin remained unchanged (Fig. S7B). As PI3K/AKT is a canonical apoptosis pathway, we also assessed Annexin-V level. However, no protective effect was evident (Fig. S7C). Likewise, Wortmannin failed to alleviate oxidative stress in aged oocytes (Fig. S7D). Hence, PI3K/AKT is unlikely to serve as a downstream effector of elamipretide in aged oocytes. To examine whether elamipretide improves oocyte quality via this pathway, we treated aged oocytes with PTC299 to inhibit VEGF-A transcriptional activity. Concentration-response analysis identified 5μM as the most effective dose for rescuing oocyte developmental quality (Fig. 9D). It was demonstrated that PTC299 reversed the age-dependent mitochondrial displacement, re-establishing the uniform peri-spindle distribution characteristic of young oocytes (Fig. 9E). In aged oocytes, VEGF-A transcriptional inhibition by PTC299 also attenuated the mitochondria-associated ROS surge (Fig. 9F, Fig. S7F). And the aberrant apoptosis triggered by oxidative stress was also ameliorated by PTC299 (Fig. 9G). Furthermore, PTC299-mediated inhibition of VEGF translation repaired the spindle-assembly defects, migration failure, and chromosome misalignment characteristic of aged oocytes (Fig. 9H). Actin filament assembly, essential for spindle migration, was likewise restored to normal levels (Fig. S7E). Together, the data position VEGF-A as a requisite node through which elamipretide restores developmental competence in aged oocytes.

**Figure 9.**
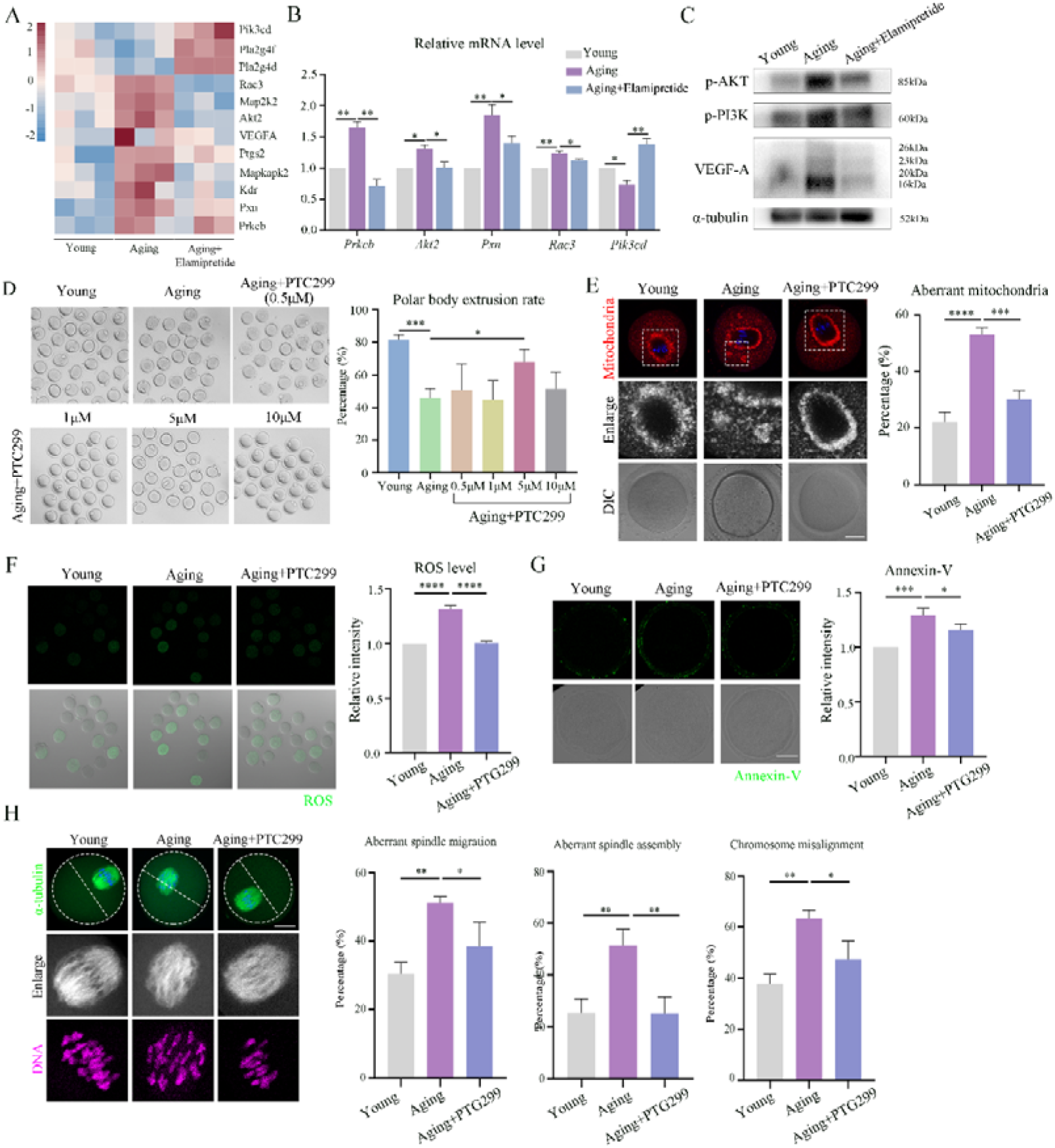
Elamipretide saves oocyte quality via VEGF-dependent pathway. (A) Heatmap of differentially expressed genes in the VEGF pathway identified by transcriptome analysis. (B) qRT-PCR was used to profile pathway gene expression, thereby confirming the reliability of the transcriptome results. (C) Western blot was used to detect the expression levels of VEGF pathway proteins in the three groups. (D) A concentration-gradient screen of VEGF inhibitors identified 5μM as the optimal treatment concentration. (E) PTC299 reversed the age-induced mis-localization of mitochondria in oocytes. Red, mitochondria; blue, DNA; scale bar, 20μm. (F) PTC299 curbed the oxidative-stress surge elicited by aging in oocytes. Green, ROS; scale bar, 20μm. (G) Inhibiting VEGF translation activity with PTC299 decreased oocyte apoptosis. Green, Annexin-V; scale bar, 20μm. (H) PTC299 restored defective spindle assembly and migration and corrected chromosome misalignment in aged oocytes. Green, α-tubulin; purple, DNA; scale bar, 20μm. * P < 0.05, ** P < 0.01, *** P < 0.001, **** P < 0.0001.

## Discussion

In present study, we proposed a novel peptide therapy method with elamipretide to reverse ovary aging for female fertility: we showed that SS31 promoted nuclear and cytoplasmic maturation of aged oocytes, showed conserved functions between mammals and *in vitro* maturation system, and elamipretide reversed age-related decline in oocyte quality through Vitamin B6-VEGF axis and ultimately improves reproductive outcomes (Figure 10).

**Figure 10.**
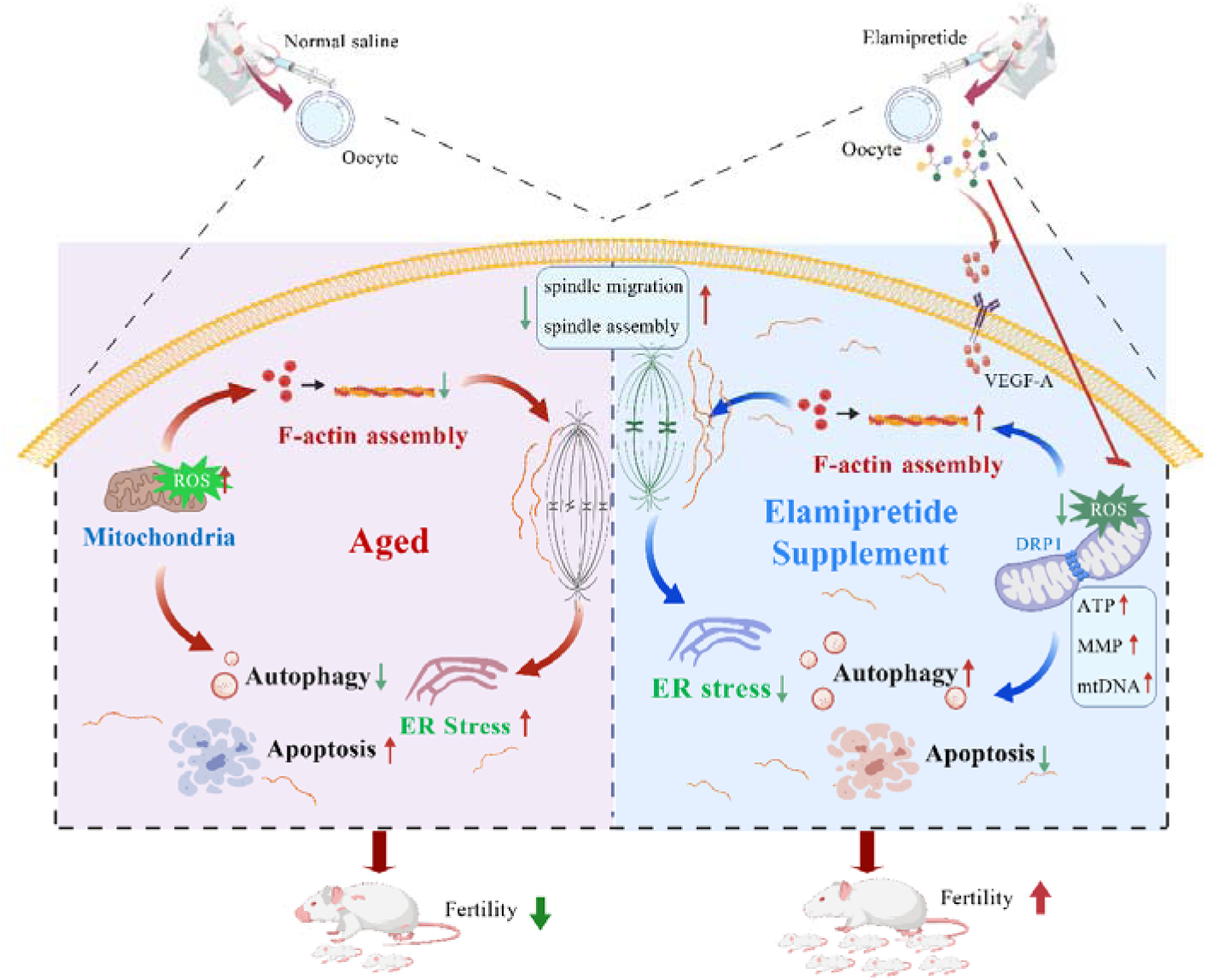
Diagram for the effects of elamipretide supplement on fertility of aged mice. SS31 promoted nuclear and cytoplasmi c maturation of aged oocytes, showed conserved functions between mammals and *in vitro* maturation system, and elamipretide reversed age-related decline in oocyte quality through Vitamin B6-VEGF axis and ultimately improves reproductive outcomes.

We first verified the therapeutic effect of elamipretide and explored the optimal treatment regimen. Previous studies have demonstrated that the injection of elamipretide into mice at concentrations of 3 mg/kg and 5 mg/kg has the best effect ^42–44^. We found that the 5mg/kg concentration of elamipretide had the best therapeutic effect on reproduction, and continuous injection of elamipretide for 7 days could effectively improve the fertility of mice aged 8, 10 months. We found that elamipretide restored the number of ovarian follicles and rejuvenated the aging ovaries, and we confirmed the restoration of the embryonic development ability of aging mice by elamipretide treatment. During porcine *in vitro* embryo production, elamipretide supplementation effectively mitigated oxidative stress damage, augmented oocytes oocyte antioxidant capacity, and markedly elevated the oocyte maturation rate and blastocyst formation rate ^45^. In aged human bone marrow-derived mesenchymal stem cells (BM-MSCs), elamipretide intervention effectively restored the functional homeostasis of mitochondria, markedly reduced intracellular ROS levels, enhanced oxygen consumption rate, and thereby reversed age-related functional decline of BM-MSCs ^46^. Additionally, elamipretide recovered the function of respiratory complexes in the aorta of aged mice, reduced the expression of inflammation-related enzymes in aged mice, and decreased elastin fragmentation ^47^.Collectively, these findings indicated that elamipretide exerted a crucial regulatory function in alleviating the aging process and enhancing oocyte quality.

We then attempted to jointly explore the potential pathways and molecular mechanisms by which elamipretide improves female fertility through metabolomics and transcriptomics analysis. In terms of overall metabolite levels, we found that elamipretide treatment could increase the overall metabolite levels of elderly female mice. Aging leads to an overall decrease in the levels of granulosa cell metabolites, especially the most significant change in pyruvate levels ^48^. We showed that the level of pyruvate significantly changed, revealing that pyruvate may be a key substance for the decline in fertility in advanced females. Pyruvate is an important component of the TCA cycle and is directly related to energy metabolism, which also suggests that changes in mitochondrial-mediated energy metabolism levels may be an important cause of infertility during female aging, which are also dated by other study ^49,50^. Our KEGG analysis also revealed for the first time that the metabolism of Vitamin B6 was significantly altered after elamipretide treatment in aging mice, suggesting that Vitamin B6 may play an important role in improving the fertility of aged mice with elamipretide. Pyridoxal 5’-phosphate (PLP) is the active form of vitamin B6 ^51^. PLP deficiency in HEK293 cells caused accumulation of lactate and pyruvate along with decreased tricarboxylic acid (TCA) cycle intermediates, thereby impairing mitochondrial oxidative metabolism ^52^. Vitamin B6 (pyridoxine) was demonstrated to synergize with vitamin B3 (nicotinamide) in regulating key molecular pathways of skeletal muscle maintenance and regeneration, thereby preventing and ameliorating age-related sarcopenia ^53^. Pyruvate has been proposed as an anti-aging metabolite that improves skin aging by generating NAD in dermal fibroblasts, thereby regulating mitochondrial and lysosome functions ^54^. We tried to directly supplement Vitamin B6, and found it can improve the oocyte quality of aging mice. Besides, our transcriptome analysis also found that there were significant differences in the overall gene expression levels between aged mice and young mice, among which the gene changes related to the TCA cycle were particularly significant, which was consistent with our metabolome results. We further conducted personalized analysis of the transcriptome results and found that elamipretide could affect the quality of oocytes in aged mice from multiple aspects such as cytoskeleton, mitochondrial function, organelle function, autophagy and apoptosis. We also screened out potential molecular pathways, VEGF pathway, by which elamipretide improves fertility in elderly female mice through KEGG and GO analyses. In vitro-cultured human umbilical vein endothelial cells (HUVECs), vitamin B6 was found to promoted AMP-activated protein kinase (AMPK) phosphorylation and vascular endothelial growth factor A (VEGF-A) synthesis ^55^. Therefore, we speculate that elamipretide regulates the metabolic process of vitamin B6 via the VEGF pathway.

We evaluated the effects of elamipretide treatment on actin filaments and microtubules in aged oocytes, which are the core drivers for the nuclear maturation of oocytes. Mitochondrial dysfunction in aged oocytes resulted in insufficient energy supply to support spindle dynamics, thereby including spindle assembly defects ^56^. In the post-ovulatory aging (POA) model of porcine oocytes, they exhibited abnormal level of cortical F-actin and acetylation of α-tubulin, which subsequently lead to spindle assembly defects ^57,58^. We found that the acetylation level of microtubules decreased and the polarity distribution of γ-tubulin was disrupted, which led to abnormal spindle morphology and chromosomal arrangement in oocytes of aging mice, and elamipretide supplementation could rescue the impairments. It was found that treatment of oocytes with MitoQ or BGP-15 reversed the spindle and chromosome abnormalities observed in reproductively aged female mice ^59^. The traditional Chinese medicine extract SCM-198 rescued aging-associated defects in oocyte spindle assembly and chromosome alignment, consequently enhancing oocyte quality ^60^. We also found that elamipretide could increase the level of cytoplasmic actin in aging oocytes. This result is consistent with another study which dated that the treatment with elamipretide promotes the rapid repair of ATP-dependent processes and also facilitates the recovery of actin cytoskeleton and cell polarity ^31^. elamipretide could affect cell function through actin, showing that elamipretide treatment reduced markers of parietal epithelial cell activation including Collagen IV, pERK1/2, and α-smooth muscle actin, and improved cytoskeletal integrity ^61^.

elamipretide is a mitochondrial-targeted peptide, and transcriptome analysis also revealed that genes enriched for inner mitochondrial membrane organization, mitochondrial metabolism, and oxidative-stress response were significantly changed after elamipretide treatment. We showed that the addition of elamipretide could restore the expression of mitochondrial related genes and mitochondrial localization, increase the mitochondrial copy number in oocytes, mediate the recovery of intracellular ATP levels and mitochondrial membrane potential, and also reduce intracellular ROS levels, indicating that elamipretide can improve mitochondrial function of aged oocytes. In a headache male mouse model, elamipretide could maintain intracellular mitochondrial homeostasis by regulating mitochondrial membrane potential through Sirt3/Pgc-1α pathway ^62^. elamipretide is also reported to be able to target cytochrome c in the mitochondrial inner membrane of damaged mitochondrial dysfunction of cardiomyocytes (CMs), and inhibits pathological ROS production ^63^. While elamipretide encompass the mitigation of oxidative stress, the suppression of inflammatory processes, and maintain mitochondrial dynamics ^64^. Treatment with elamipretide improved mitochondrial respiratory capacity and promoted supercomplex organization, which benefits on improving cardiac mitochondrial dysfunction in the model of Barth Syndrome mice ^65^. Therefore, the mitochondria-targeted peptide elamipretide can significantly improve the maturation quality of aged oocytes by targeting and regulating mitochondrial function.

Normal organelle function is a fundamental prerequisite for the cytoplasmic maturation of oocytes. We found that elamipretide can restore the expression and localization of Golgi apparatus, and the functions showing by Rab10, Rab11, and GM130, and elamipretide induced a decrease in ER stress level mediated by GRP78, restored the normal distribution of lysosomes and reduced the level of autophagy in aged oocytes. We indicate that elamipretide treatment is capable of restoring Golgi apparatus function and reducing ER stress in aged oocytes. The treatment with elamipretide can inhibit the release of cytokines in skeletal muscle cells mediated by the Golgi apparatus ^66^, and treatment with elamipretide can reduce ER stress in leukocytes from type 2 diabetes in patients ^67^. elamipretide mediates the regulation of autophagy levels by regulating cPLA2-induced lysosomal membrane permeability in rat ^68^. elamipretide promotes the formation of autolysosomes and autophagosomes, thereby facilitating autophagic flux to a certain degree in human hepatocellular carcinomas ^69^. Besides, elamipretide was reported to promote PHB2-mediated mitophagy activation to inhibit mtDNA release in mouse microglial cell (BV2) ^70^. Thus, elamipretide is able to effectively ameliorate oocyte quality by repairing organelle dysfunction in aged oocytes.

As *in vitro* maturation of oocytes is widely used by assisted reproductive technology (ART), we explored the effects of elamipretide on *in vitro* matured oocytes. Similar with *in vivo* situation, supplement with elamipretide in the culture medium also improved the maternal aged mouse oocyte and post-ovulatory porcine oocyte maturation. And elamipretide also could decrease oxidative-stress level, reverse the mitochondria dysfunction and rescue the abnormally high apoptosis back to normal levels. Similarly, the supplementation of elamipretide could improve the quality of porcine early embryos ^45^. Taken together, the results demonstrate that elamipretidecan directly target oocytes and embryos at the cellular level to improve their development potential during *in vitro* culture.

Subsequently, we tried to explore the molecular mechanisms underlying the improvement of female fertility by elamipretide. By integrating metabolomic and transcriptomic data, we identified the VEGF pathway as a key target. We showed that elamipretide supplementation restored the expression levels of VEGF-A and its downstream PI3K/AKT proteins. Treatment with PTC299 to inhibit VEGF-A transcriptional activity on aged oocytes showed similar effects with elamipretide, and improved the quality of aging oocytes. VEGF served as a pivotal upstream factor that activates the PI3K/AKT pathway in endothelial cells, and the VEGF-VEGFR2-PI3K/AKT signaling axis constitutes a key molecular mechanism regulating angiogenesis and vascular homeostasis maintenance ^71^. elamipretide has been demonstrated to conjugate with anti-VEGFR2 and exert therapeutic effects in diabetic nephropathy (DN) models by reducing pro-inflammatory factors and fibrosis markers, as well as alleviating oxidative stress ^72^. In aortic stenosis (AS) treatment, VEGF protects cardiac function in myocardial infarction models via the canonical VEGF-PI3K-AKT signaling pathway as well as the mitochondrial anti-apoptotic pathway ^73^. A conductive hydrogel was found to enhance the angiogenic capacity of vascular endothelial cells under senescent conditions via VEGF/ VEGFR2 upregulation and the PI3K-AKT-eNOS pathway activation ^74^. However, we next tested whether elamipretide affects female fertility through PI3K/AKT pathway and found that the expression levels of PI3K/AKT pathway-related proteins in aged oocytes did not change significantly after the treatment of elamipretide. Besides, wortmannin failed to alleviate oxidative stress in aged oocytes. Hence, PI3K/AKT is unlikely to serve as a downstream effector of elamipretide in aged oocytes. These results dated that elamipretide improved oocyte quality of aged mice through VEGF pathway.

In summary, our study showed that short-term treatment with elamipretide can improve the fertility of aged female, and it is mainly through effects on the rescue of cytoplasmic and nuclear maturation of oocytes by vitamin B6-based VEGF pathway.

## Ethical Approval and Consent to Participate

All use procedures of animals and experimental programs were carried out in accordance with the guidelines of the Animal Research Ethics Committee of Nanjing Agricultural University and approved by the Animal Research Ethics Committee. Human oocyte collection was followed to the guideline of ethic committee of Maternity and Child Health Care of Guangxi Zhuang Autonomous Region, and the patients were aware and permitted the use of the oocytes.

## Consent for publication

All authors are consent for the publication of this study.

## Availability of data and material

All data generated or analyzed during this study are included in this published article.

## Authors’ contributions

HLZ, SCS designed the study; HLZ, YW, CW, XG performed the experiments; HC, YXH, XW, XTY, ZJW, WLP. RJM, PSL, JS contributed the materials; HLZ, YW, SCS wrote the manuscript; HLZ, YW, SCS analyzed the data. All the authors approved the final manuscript.

