## Supplemental Materials for "Elamipretide reverses female fertility decline during reproductive aging via regulating VEGF in oocytes"

**Running Title:** Elamipretide for aging-induced infertility

^#^These authors contributed equally to this work.

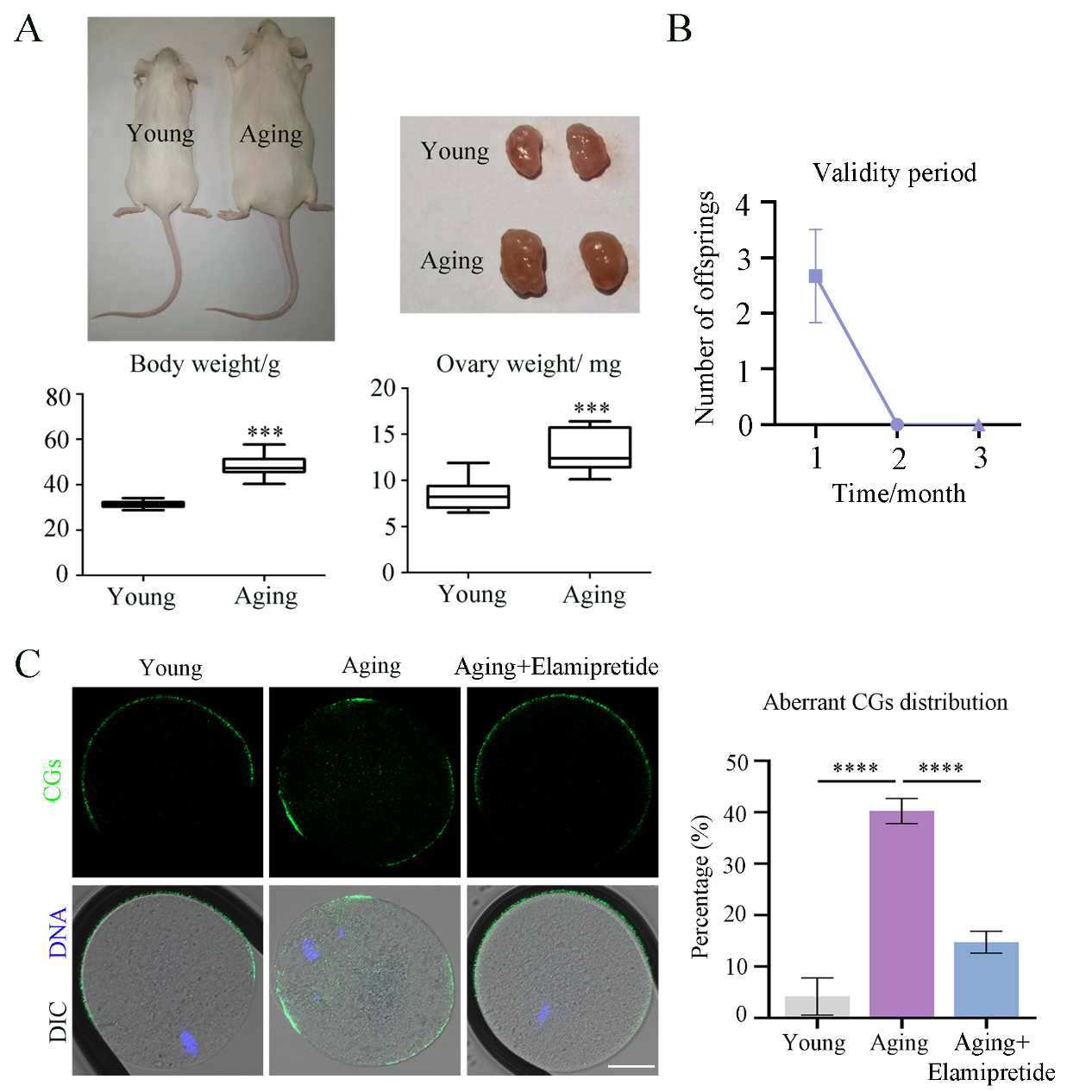
**Figure S1.** Elamipretide orchestrates cortical-granule trafficking in aged mouse oocytes. (A) Comparison of body and ovarian weights between aged and young mice. (B) The restorative effect of elamipretide on litter size is transient. (C) elamipretide reversed aging-induced defects in cortical granule trafficking. Green, cortical granule; blue, DNA; scale bar, 20μm. *** P < 0.001, **** P < 0.0001.

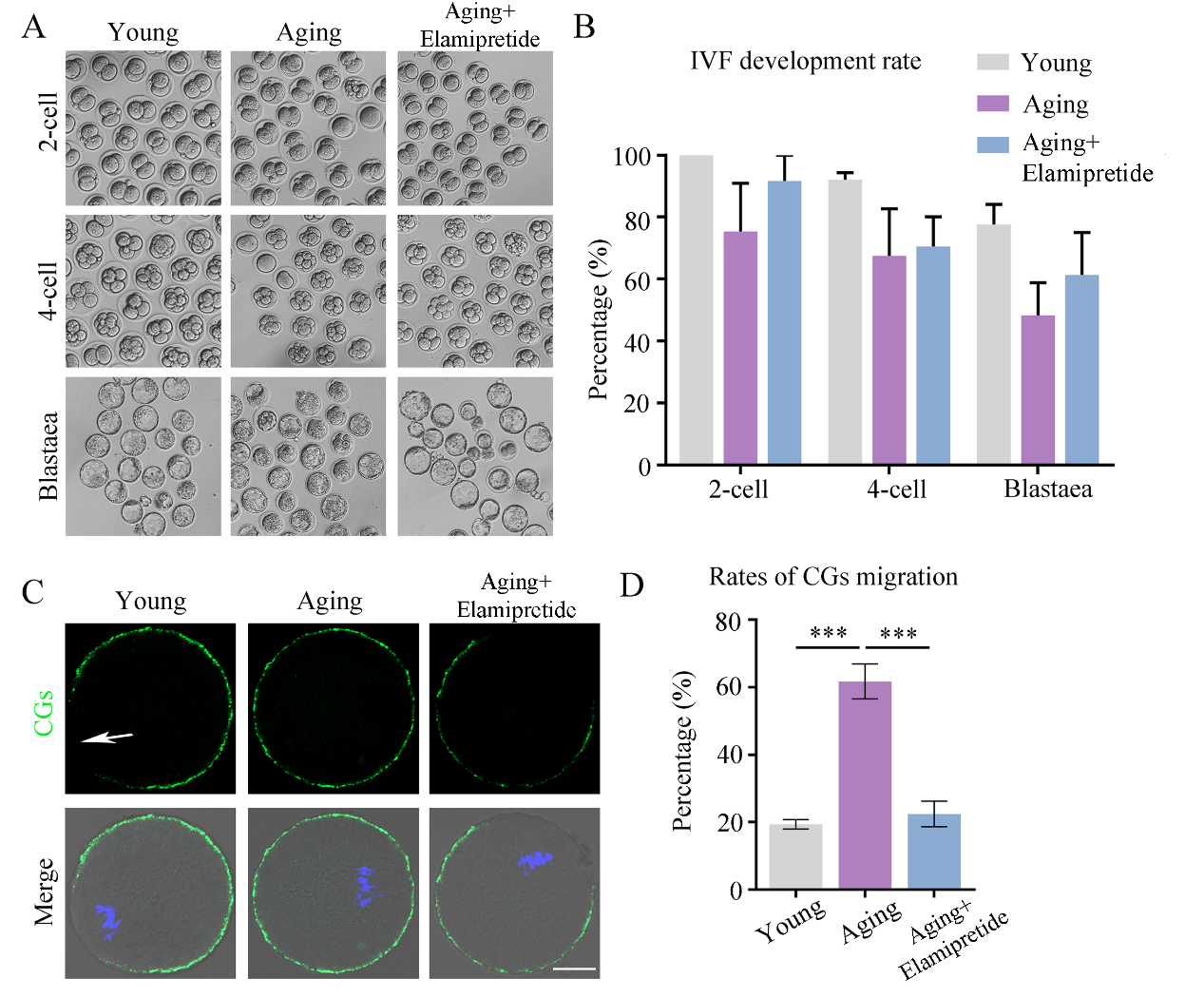
**Figure S2.** In vitro elamipretide improves embryo quality. (A) In vitro elamipretide partially rescued IVF embryo developmental quality. (B) The statistical analysis of IVF embryo development quality. (C) In vitro elamipretide supplementation accelerated CGs migration in aged oocytes. Green, cortical granule; blue, DNA; scale bar, 20μm. (D) The statistical analysis of the rate of CGs migration. *** P < 0.001, **** P < 0.0001.

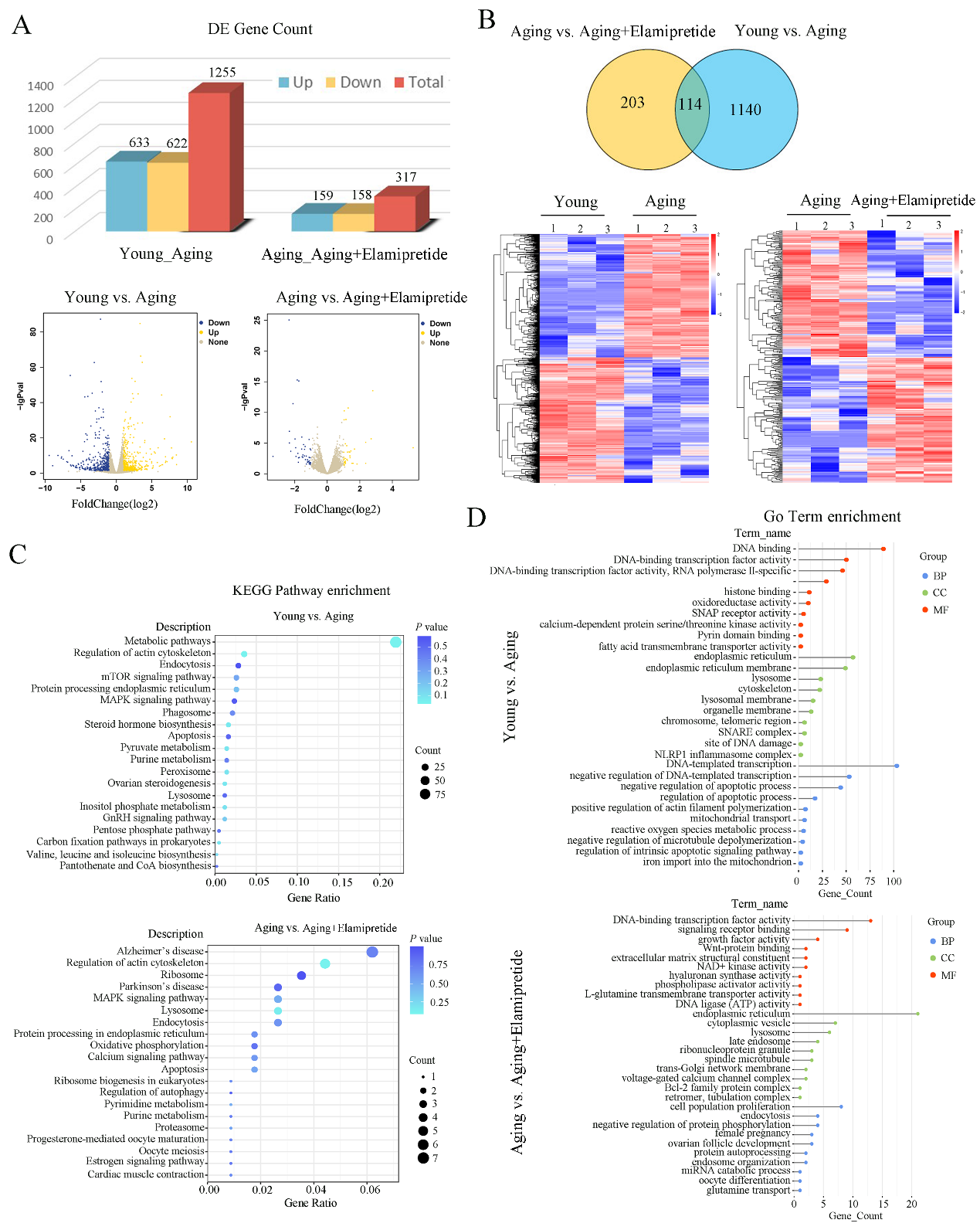
**Figure S3.** Transcriptome profiling of oocytes after in vitro elamipretide supplementation. (A) Statistics of up- and down-regulated DEGs in aging vs. young and aging+elamipretide vs. aging oocytes. (B) Venn diagram and heatmap clustering analysis of differentially expressed genes. (C) KEGG pathway enrichment clustering of differentially expressed genes. (D) GO term enrichment clustering analysis of differentially expressed genes.

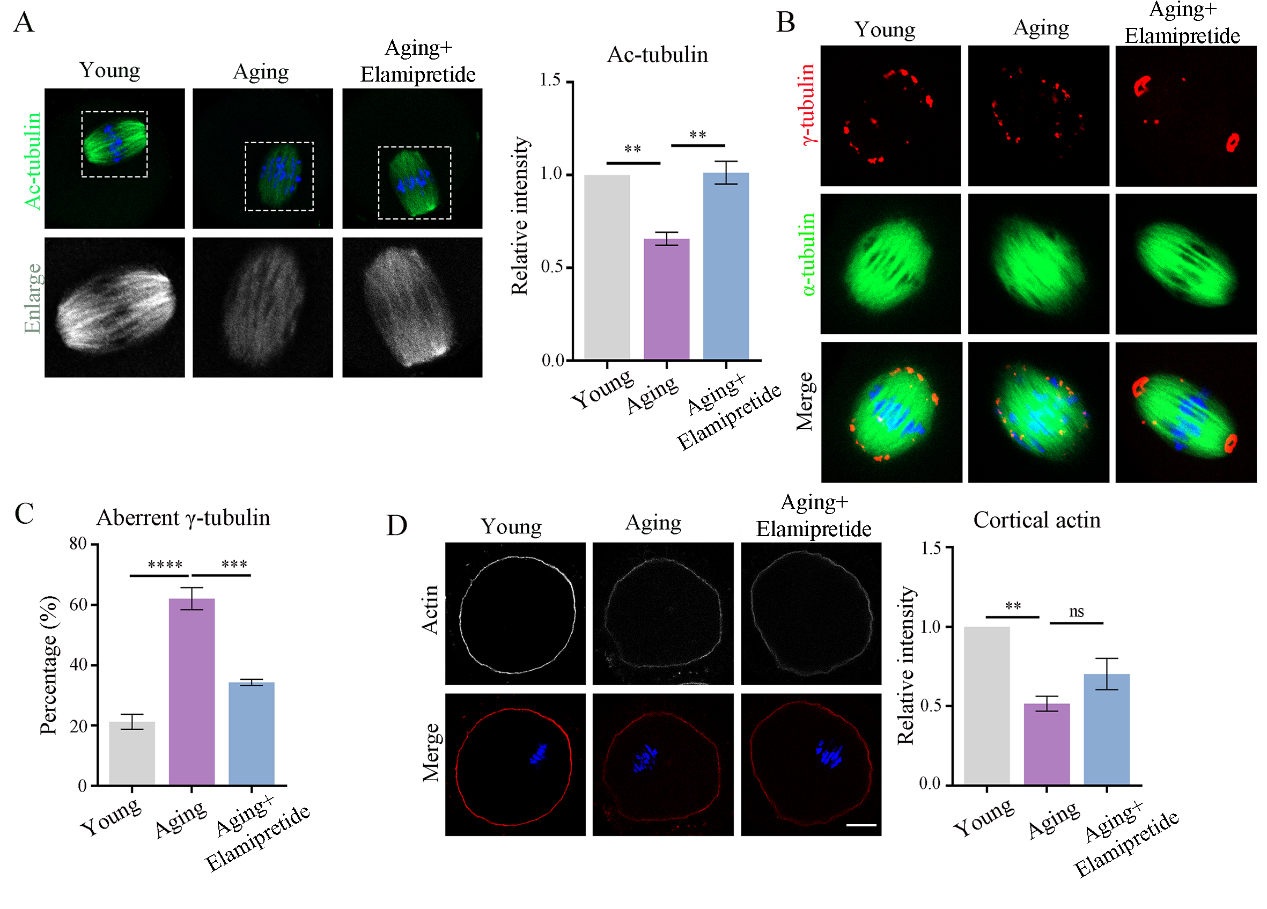
**Figure S4.** In vitro elamipretide supplementation restores cytoskeletal assembly defects in aged oocytes. (A) elamipretide elevated microtubule acetylation and steadied the cytoskeleton in aged oocytes. Green, ac-tubulin; blue, DNA; scale bar, 20μm. (B) elamipretide supplementation restored γ-tubulin distribution in aged oocytes. Green, α-tubulin; red, γ-tubulin; blue, DNA; scale bar, 20μm. (C) The statistical analysis of the rate of aberrant γ-tubulin. (D) elamipretide supplementation only partially rescued cortical actin fluorescence intensity in aged oocytes. Red, actin; blue, DNA; scale bar, 20μm. ** P < 0.01, *** P < 0.001, **** P < 0.0001, and ns, no significant difference.

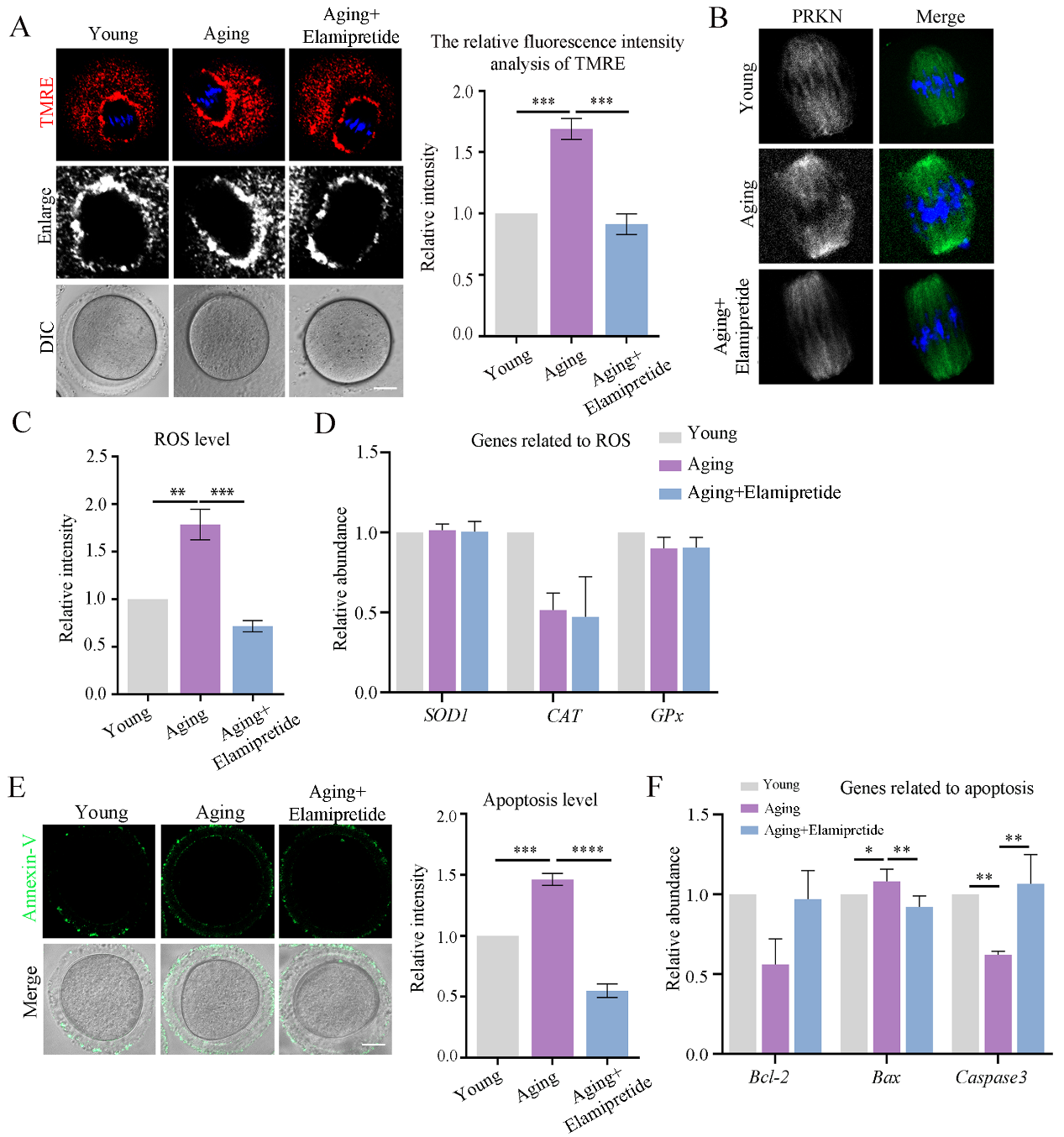
**Figure S5.** In vitro elamipretide supplementation improves mitochondrial function in aged oocytes. (A) elamipretide supplementation alleviated the abnormally elevated membrane potential of aged oocytes. Red, TMRE; blue, DNA; scale bar, 20μm. (B) elamipretide intervention rescued the increase of mitophagy protein Parkin exhibited by aged oocytes. Green, Parkin; blue, DNA; scale bar, 20μm. (C) Statistics showing the reverse of ROS levels by in vitro elamipretide supplementation. (D) elamipretide supplementation failed to restore SOD1, CAT, and GPx activity. (E) elamipretide intervention significantly reduced apoptosis in aged oocytes. Green, Annexin-V; scale bar, 20μm. (F) elamipretide treatment in vitro normalized the expression of apoptosis-linked genes. * P < 0.05, ** P < 0.01, *** P < 0.001, **** P < 0.0001, and ns, no significant difference.

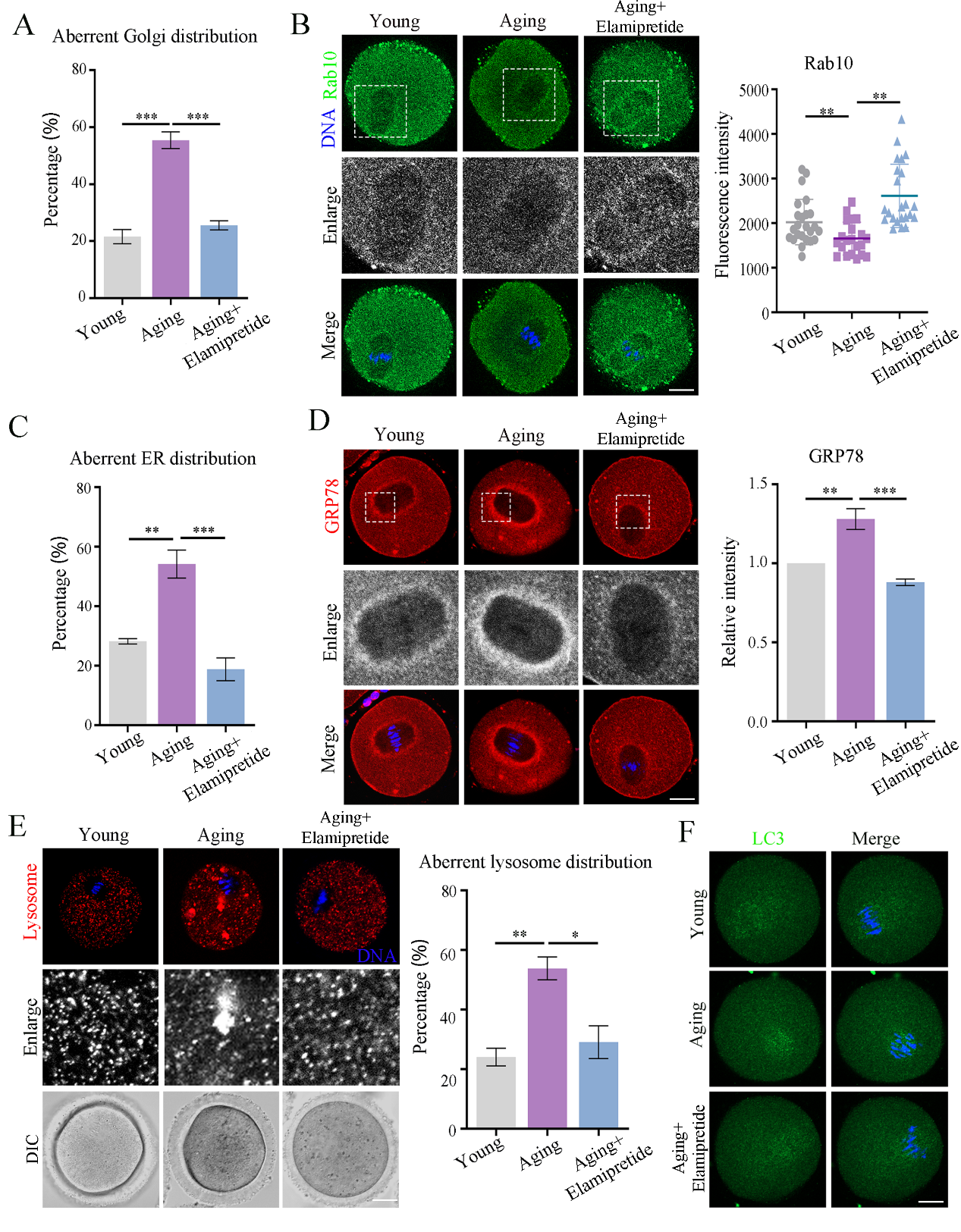
**Figure S6.** Elamipretide resets organelle layout and performance in aged oocytes. (A) The statistical analysis of aberrant distribution of Golgi after elamipretide supplementation in aged oocytes. (B) elamipretide supplementation elevated Rab10 expression in aged oocytes. Green, Rab10; blue, DNA; scale bar, 20μm. (C) Statistical analysis of ER distribution upon in vitro elamipretide supplementation. (D) In vitro elamipretide supplementation alleviated ER stress in aged oocytes. Red, GRP78; blue, DNA; scale bar, 20μm. (E) Exogenous elamipretide supply dispersed excessive lysosomal clustering in aged oocytes. Red, lysosome; blue, DNA; scale bar, 20μm. (F) elamipretide supplementation significantly reduced the number of LC3 autophagic vesicles in age oocytes. Green, LC3; blue, DNA; scale bar, 20μm. * P < 0.05, ** P < 0.01, *** P < 0.001.

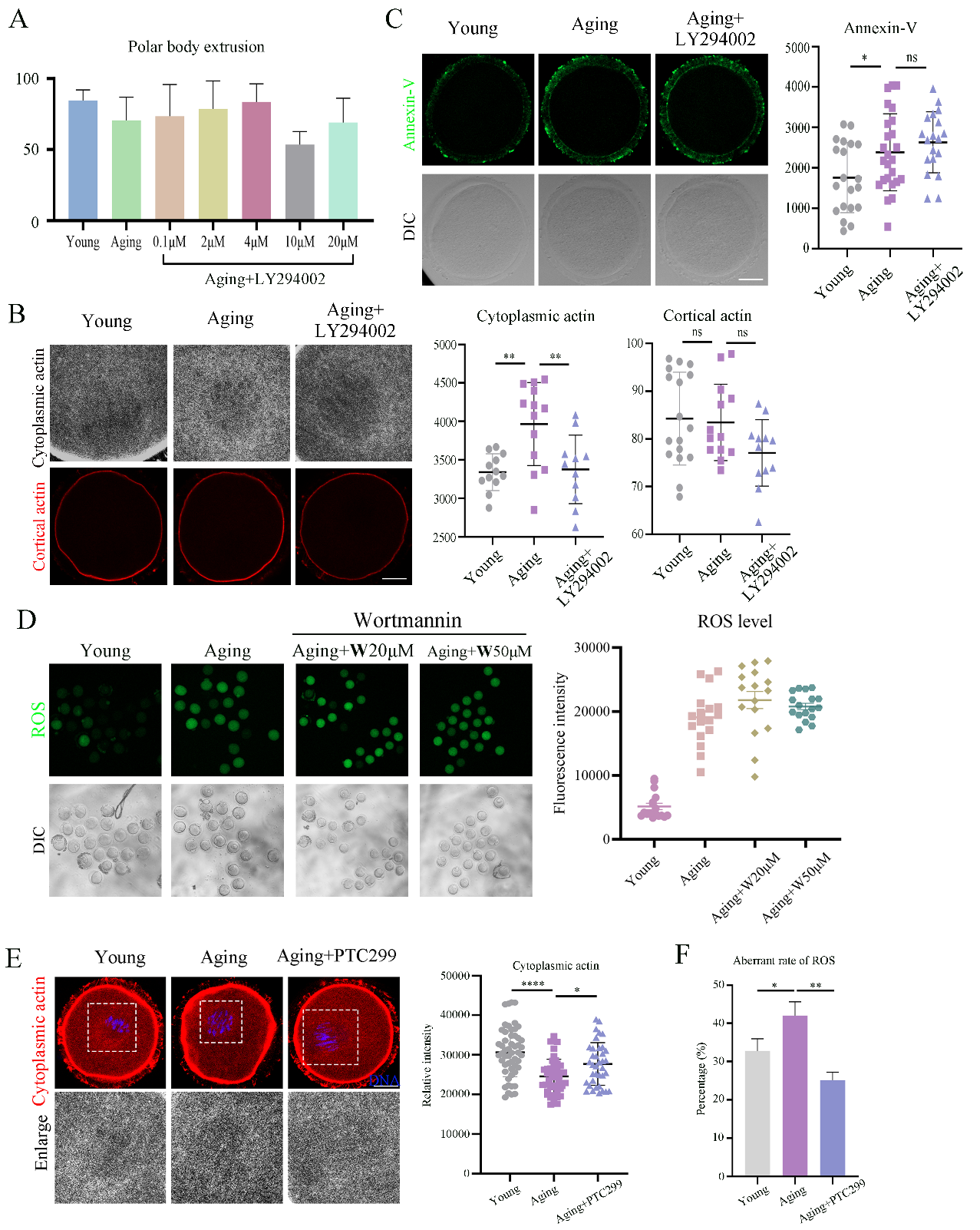
**Figure S7.** Elamipretide fails to modulate aged oocyte quality through the PI3K/AKT pathway. (A) PI3K/AKT inhibitor LY294002 only partially restored aged oocytes maturation. (B) LY294002 treatment recovered the abnormally elevated cytoplasmic actin levels in aged oocytes. Grey, cytoplasmic actin; red, cortical actin; scale bar, 20μm. (C) LY294002 failed to mitigate the heightened apoptosis observed in aged oocytes. (D) Another PI3K/AKT inhibitor Wortmannin failed to reverse ROS levels in aged oocytes. Green, ROS; scale bar, 20μm. (E) The VEGF inhibitor PTC299 reversed the decline in cytoplasmic actin levels in age oocytes. Red, actin; blue, DNA; scale bar, 20μm. (F) The statistical analysis of PTC299 reversing ROS levels in aged oocytes. * P < 0.05, ** P < 0.01, **** P < 0.0001, and ns, no significant difference.

**Table S1. Primers used in the study.**

| Drp1 | F 5'-CAGGTGGTGGGATTGGAGAC | R 5'-CTGGCATAATTGGAATTGGTTT |
| --- | --- | --- |
| Opa1 | F 5'-CCGAGGATAGCTTGAGGGTT | R 5'-CGTTCTTGGTTTCGTTGTGA |
| PGC1-α | F 5'-TATGGAGTGACATAGAGTGTGCT | R 5'-CCACTTCAATCCACCCAGAAAG |
| ATP5B | F 5'-GGTTCATCCTGCCAGAGACTA | R 5'-AATCCCTCATCGAACTGGACG |
| SOD2 | F 5'-CAGACCTGCCTTACGACTATGG | R 5'-CTCGGTGGCGTTGAGATTGTT |
| Gpx3 | F 5'-CCTTTTAAGCAGTATGCAGGCA | R 5'-CAAGCCAAATGGCCCAAGTT |
| Ndufs3 | F 5'-TGGCAGCACGTAAGAAGGG | R 5'-CTTGGGTAAGATTTCAGCCACAT |
| Caspase3 | F 5'-ATGGAGAACAACAAAACCTCAGT | R 5'-TTGCTCCCATGTATGGTCTTTAC |
| Jun | F 5'-CCTTCTACGACGATGCCCTC | R 5'-GGTTCAAGGTCATGCTCTGTTT |
| BAX | F 5'-TGAAGACAGGGGCCTTTTTG | R 5'-AATTCGCCGGAGACACTCG |
| PARP1 | F 5'-GGCAGCCTGATGTTGAGGT | R 5'-GCGTACTCCGCTAAAAAGTCAC |
| Capn1 | F 5'-ATGACAGAGGAGTTAATCACCCC | R 5'-GGCTATGAGAAACCGGAGGG |
| GM130 | F 5'-TGTCACAGAACCGAGAGCTGA | R 5'-GCTGCTCCTGTAGACCCTG |
| Rab11 | F 5'- CTCTGGACGAGGTCTTCCG | R 5'-TGTTCCGTGTGAACTGGATGG |
| GRP78 | F 5'-ACTTGGGGACCACCTATTCCT | R 5'-ATCGCCAATCAGACGCTCC |
| ERO1L | F 5'-TTCTGCCAGGTTAGTGGTTACC | R 5'-GTTTGACGGCACAGTCTCTTC |
| CHOP | F 5'-CTGGAAGCCTGGTATGAGGAT | R 5'-CAGGGTCAAGAGTAGTGAAGGT |
| ATF4 | F 5'-ATGGCGCTCTTCACGAAATC | R 5'-ACTGGTCGAAGGGGTCATCAA |
| LC3 | F 5'-GACCGCTGTAAGGAGGTGC | R 5'-CTTGACCAACTCGCTCATGTTA |
| ATG7 | F 5'-GTTCGCCCCCTTTAATAGTGC | R 5'-TGAACTCCAACGTCAAGCGG |
| NRBF2 | F 5'-AAGGACCCCTCAACCTTGCT | R 5'-CAGTTCCAGTGATAAGTGAGCC |
| LAMP2 | F 5'-TGTATTTGGCTAATGGCTCAGC | R 5'-TATGGGCACAAGGAAGTTGTC |
| Prkcb | F 5'-AAGGAGCATGCGTTTTTCCG | R 5’-GTCTCGCTTGTCTCTAGCTTTTG |
| Akt2 | F 5’-CGCCGGTGACAGACGATA | R 5’-TTTGTGGAGCCAGCCTTCTT |
| Pxn | F 5’-GACGACCTCGATGCCCTG | R 5’-GGGGCTCCTCTGACAAGAAC |
| Rac3 | F 5’-TATCCCCACAGTTTTCGACAAC | R 5’-GAGAGTGGCCGAAGCCTAT |
| Pik3cd | F 5’-TTAATCTCCCAGGCAGAGGG | R 5’-CTGAGGTCTGTTTTCCTGTTTGG |
